# Ancestral phototrophic *Rhizobiaceae* evolved in association with algae, then plants

**DOI:** 10.64898/2026.08.14.744861

**Authors:** Steven B. Kuzyk, Philipp Halama, Mohit Kumar Saini, Mathias Müsken, Michal Koblížek, Jörg Overmann

**Affiliations:** Department of Microbial Ecology and Diversity Research, Leibniz-Institute DSMZ-German Collection of Microorganisms and Cell Cultures, Inhoffenstraße 7B, 38124 Braunschweig, Germany; Laboratory of Anoxygenic Phototrophs, Institute of Microbiology CAS, 37901 Třeboň, Czechia; Central Facility for Microscopy, HZI - Helmholtz Centre for Infection Research; Inhoffenstraße 7, 38124 Braunschweig, Germany; Braunschweig University of Technology, Universitätspl. 2, 38106 Braunschweig, Germany; Bavarian State Collections of Natural History (SNSB), Menzinger Straße 71, 80638 Munich, Germany; Chair for Molecular Diversity Research, Faculty of Biology, Ludwig Maximilian University of Munich, Menzinger Straße 71, 80638 Munich, Germany

**Author notes:** Corresponding author. Address: Inhoffenstraße 7B, 38124 Braunschweig, Germany. Corresponding author. Address: Menzinger Straße 71, 80638 Munich, Germany.

**Keywords:** Microbial symbiosis, Rhizobia, Aerobic anoxygenic phototrophs, Microbial consortia, Syntrophy, Phycosphere, Microbial evolution, Bacteriochlorophyll, *Hoeflea*, *Peteryoungia*

## Abstract

*Rhizobiaceae* serve as classical models for elucidating mutualistic plant-microbe interactions yet they represent a narrow phylogenetic subgroup of *Alphaproteobacteria*. Studying additional lineages of *Rhizobiaceae*, we observed broad associations with oxygenic phototrophs beyond land plants, including early branching clades of submerged plants, multicellular and unicellular algae, as well as cyanobacteria. In particular, bacteria of the genus *Hoeflea* were often affiliated with cyanobacteria or microbial algae, whereas *Peteryoungia* spp. colonized roots of submerged plants. While both genera were originally described as nonpigmented heterotrophs, our detailed genomic, biochemical and physiological analyses revealed that most strains actually contained genes for anoxygenic photosynthesis. Under oligotrophic, oxic growth conditions, each characterized representative expressed bacteriochlorophyll *a*-containing functional photosynthetic complexes. Photosynthesis genes shared the highest homology among phylogenetically closest relatives, displaying topologies congruent to cognate house-keeping gene phylogenies, and maintained highly conserved gene synteny across the chromosomes of different species. Together, this indicated a vertical inheritance and long ancestral history of aerobic anoxygenic photosynthesis in *Rhizobiaceae* rather than multiple recent horizontal transfers. Subsequent time-scale phylogenetic analysis suggested that the last common ancestor of *Rhizobiaceae* was an aquatic phototroph, with different lineages of *Rhizobiaceae* consecutively evolving in association with algae, land plants, then later legumes. While aquatic lineages maintained photosynthetic machinery till today, *Rhizobia* which developed symbioses with land plants either as mutualistic endosymbiosis within root nodules or as plant pathogens, concomitantly lost photosynthetic capability. Based on our results, multiple biotic interactions with diverse oxygenic phototrophs drove the early evolution of *Rhizobiaceae*.

## Introduction

Rhizobia have largely been recognized as the typical nitrogen-fixing bacterial symbionts of plants [**Editorial Board, 1969; Editorial Secretary, 1970**], where signal exchange with hosts enable root-hair colonization and formation of bacteroid nodules [**Poole *et al.,* 2018**]. As endosymbionts, they generate excess fixed nitrogen to support plant growth in exchange for fixed organic carbon [**Zhang *et al.,* 2024**]. Diverse *Rhizobiaceae*, such as *Rhizobium leguminosarum* or *Sinorhizobium meliloti*, develop mutualistic partnerships with distinct host species [**Martinez-Romero *et al.,* 2024**], while the closely related *Agrobacterium* spp. are pathogens causing plant diseases in different plants [**Flores-Félix *et al.,* 2020**]. Host specificity is determined genetically [**Menna and Hungria, 2011; Wardell *et al.,* 2021**], where specialized cell signaling mediated by nodulation (*nod*) or virulence (*vir*) factors are required for each association.

Although *Rhizobiales* bacteria of the genus *Hoeflea* were originally defined as marine, non-nodulating, aerobic heterotrophs of based on the type species *H. marina* [**Peix *et al.,* 2005**], subsequent isolation of *H. phototrophica* from the marine dinoflagellates *Alexandrium lusitanicum* and *Prorocentrum lima* suggested a link with algae [**Biebl *et al.,* 2006**]. The later cultivation of *H. alexandrii* from *Alexandrium minutum* [**Palacios *et al.,* 2006**], *H. anabaenae* from the cyanobacterium *Anabaena* [**Stevenson *et al.,* 2011**], *H. prorocentri* from *Prorocentrum mexicanum* [**Yang *et al.,* 2018**], and both *H. algae* and *H. ulvae* from marine macroalgae [**Baek *et al.,* 2023**] further support a *Hoeflea* relationship with algae. Recently, metagenomic and isotopic analysis of yet another lineage of the *Rhizobiales* (‘*Candidatus* Tectiglobus diatomicolá) revealed a tight endosymbiotic partnership with a marine diatom [**Tschitschko *et al.,* 2024**]. These observations suggest that the capability to associate with oxygenic phototrophs evolved in many more lineages of the *Rhizobiales* than previously recognized and also extends towards multicellular and unicellular algae, or cyanobacteria as hosts.

Beyond the production of various (or absent) *nod* factors to initiate host-specific nodulation, members of the *Rhizobiaceae* feature considerable physiological heterogeneity, with broad metabolic capabilities, transformations of toxic metal(loids) [**Deng, 2020**], and non-canonical nitrogen fixation [**Okazaki *et al.,* 2015**]. One unusual, so far rare trait is the production of bacteriochlorophyll (BChl) *a* and the capability of anoxygenic photosynthesis, first detected in *Bradyrhizobium* spp. [**Evans *et al.,* 1990; Giraud *et al.,* 2000**]. While both photosynthetic and non-photosynthetic strains of this genus are capable of nodulating *Aeschynomene* without *nod* factors [**Fleischman and Kramer, 1998; Camuel *et al.,*2023**], those with photosynthesis (such as BTAi1 or ORS278) show increased nitrogen fixation rates and significantly promote the growth of their host [**Giraud *et al.,* 2000**]. Notably, *Hoeflea phototrophica* was later shown to also be an aerobic anoxygenic phototrophic (AAP) bacterium, harvesting light by BChl *a* under aerobic conditions [**Biebl *et al.,* 2006**]. Till now, the lack of other photosynthetic representatives of the *Rhizobiales* has largely prevented an analysis of the evolutionary origin and inheritance [**Fleischman and Kramer 1998; Avontuur *et al.,* 2023**], or adaptive value of anoxygenic photosynthesis in this ecologically relevant group [**Camuel *et al.,* 2023**]. Conversely, AAPs have been associated with algae since their initial discovery [**Harashima *et al.,* 1978**], often correlated with phytoplankton blooms worldwide [**Koblížek, 2015; Gazulla *et al.,* 2024; Cai *et al.,* 2024; Kust *et al.,* 2025**], frequently isolated from the phycosphere [**Yurkov and Beatty 1998**; **Tinguely *et al.,* 2023; Nedaschkovskaya *et al.,* 2025**], and even found directly attached to algae [**Wagner-Dobler *et al.,* 2010; Kuzyk *et al.,* 2022; Tanabe *et al.,* 2023**].

We conducted a systematic survey of phototrophic capacity in all described species of the *Rhizobiaceae*. While most taxa were previously considered as unpigmented strict heterotrophs, we specifically targeted *Hoeflea* spp. alongside unexpected *Peteryoungia* spp. since our preliminary analyses of preexisting draft genomes had indicated the presence of the necessary genes. Combined physiological, biophysical, and phylogenetic approaches allowed us to track the *in vivo* functionality and distribution of photosynthesis genes across all *Rhizobiaceae*, permiting evolutionary origin and overarching ecological implications to be elucidated.

## Materials and Methods

### Strains utilized

*Hoeflea marina* DSM 16791^T^*, Hoeflea alexandrii* DSM 16655^T^*, Hoeflea phototrophica* DSM 17068^T^*, Hoeflea siderophila* DSM 21587^T^*, Hoeflea suaedae* DSM 23348^T^, *Peteryoungia rosettiformans* DSM 26376^T^, *Peteryoungia aggregata* DSM 1111^T^ and *Peteryoungia rhizophila* DSM 103161^T^ were reactivated from the DSMZ culture collection. *Hoeflea olei* KCTC 42071^T^, *Peteryoungia ipomoeae* LMG 27163^T^, *Peteryoungia desertarenae* JCM 33657^T^ and *Dunaliella bicculata* SAG 19-4 were obtained from other respective culture collections (Table S1). All strains were accessed in compliance with Nagoya Protocol requirements [**Faggionato et al., 2026**], as stated in the Acknowledgements. *H. alexandrii* DSM 118613 was isolated from a *Oxyrrhis marina* SAG 21.89 culture fed with *Dunaliella bicculata* SAG 19-4, while *H. anabaenae* WH2K was reisolated from *Anabaena variabilis* SSM-00 consortia provided by Dr. John B. Waterbury. All strains were grown at 18°C at 20 µmol photons m^-2^s^-1^ light with a combination of three flourescent bulbs (Osram L30W/830 Lumilux warm white, Osram L30W/840 Lumilux cool white, Osram L30W/77 Fluora) in 18h:6h light:dark cycles. Every *Rhizobiales* spp. was cultured on PY+ medium at pH 7.0, a version of PY [**Stevenson et al., 2011**] with modifications (Table S2). Briefly, in addition to diluted yeast extract and peptone, it comprised an artificial sea water (ASW) solution modified from DSMZ S-3501 [**Koblitz et al., 2022**], with trace elements and vitamin solutions designed for AAP [**Yurkov et al., 1999**]. *A. variabilis* grew on ½ SO medium [**Waterbury et al., 1986**] with modifications (Table S2), while *D. bicculata* was propagated with f/2 medium [**Borowitzka and Siva, 2007**; DSMZ m1877].

Co-cultures of *H. anabaenae* with *A. variabilis*, and *H. alexandrii* DSM 118613 with *D. bicculata* were imaged via field-emission scanning electron microscopy (SEM) using a Zeiss Merlin (Oberkochen, Germany) as described with minor modifications [**Viera *et al.,* 2020**]. In brief, 2% glutaraldehyde or 2% gluteradehyde + 5% formaldehyde were utlized to fix either consortium, respectively, with both critical-point dried under liquid CO_2_ (CPD 300, Leica Microsystems, Wetzlar) during processing.

### Growth kinetics

Five *Hoeflea* and five *Peteryoungia* species were chosen for anoxygenic photosynthesis induction assays (Table S1). Triplicates of 5 mL test tubes were inoculated with 5% (v/v) cultures and incubated at 20°C in the dark, as this promotes strongest AAP pigmentation [**Kuzyk *et al.,* 2023**]. Biomass was monitored via optical density (OD) measurements at 660 nm using a Genesys 20 spectrophotometer (ThermoScientific, Braunschweig, Germany) on days 0 and 14. Salinity ranges were determined by modifying PY+ medium with 0, 0.5, 1, 2, 3, 4, 5, 7.5, 10, or 15× ASW. To test the impact of vitamins and trace elements on growth and pigmentation, PY+ medium without trace elements, vitamins, or neither, was utilized and compared to the full PY+ medium for all 10 strains. Effects of nutrient load on cell pigmentation were determined by adjusting the complex organics (bactopeptone and yeast-extract in a 6:1 (w/w) ratio) to higher concentrations [g/L] of 0, 0.35, 1.75, 3.5, 7.0, 10.5, 21.0, 31.5, 70, 105, keeping trace vitamins, metals, and salinity constant as in the original PY+ recipe. After cultivation, an aliquot of each sample was collected to determine BChl *a* production levels.

### Spectroscopy, pigment analysis, and photosynthetic activity

Whole-cells were resuspended in 0.3 mL 20 mM pH 7.8 Tris-HCl buffer and 0.7 mL glycerol to reduce light scattering, prior to spectroscopic measurement [**Kuzyk *et al.,* 2023**]. Absorption spectra of cells or extracts were recorded using a Shimadzu UV2600 equipped with an integrating sphere (Shimadzu, Kyoto, Japan), or a UV-Vis Lambda 365+ spectrophotometer (Perkin Elmer, Weiterstadt, Germany) from 300 to 1100 nm in 10 mm path-length quartz cuvettes. Pigments were extracted from pelleted cells (10,000 ×*g* for 5 min) using 0.5 mL acetone:methanol (7:2), cellular debris were removed via centrifugation (10,000 ×*g* for 5 min), and pigments analyzed by direct spectrophotometry or separately via high-performance liquid chromatography (HPLC). BChl *a* concentrations were determined at the peak absorbance wavelength of 769 nm in solvent extracts quantified via standard curves of purchased pigments (Sigma Aldrich, Germany) [**Kuzyk *et al.,* 2023**]. HPLC permitted pigment identification by injecting extracts into Shimadzu Nexera system equipped with a Kinetex C8 column [**Nupur *et al.,* 2021**], followed by absorption spectra of each eluted product via a diode-array detector, and comparison to an internal library with bacterial pigment standards. Suspensions of bacterial cells were also subjected to fluorescence induction-decay kinetics to determine photochemical yield and to confirm photosynthetic activity *in vivo* [**Koblížek *et al.,* 2010; Kaftan *et al.,* 2019**].

### Genome sequencing and annotations

DNA was extracted from 12 type strains (Table S3), with fragmentation sizes quantified via FemtoPulse (Agilent, CA, USA), where DNA of sufficient mean lengths >6000 bp was either sequenced via PacBio Sequel *IIe* as described before [**Methner *et al.,* 2023**], or via a two platform approach: Oxford Nanopore long-reads mapped with high-quality Illumina NextSeq 2000 paired-end sequences of 2×150 bp as described [**Kuzyk *et al.,* 2022**], depending on equipment availability. Long-read sequences generated by PacBio were assembled using Flye v2.9 [**Kolmogorov *et al.,* 2019**], while short Illumina reads were mapped to the NanoPore assembly using Minimap2 [**Li, 2018**]. All chromosomes were circularized and rotated to start locus tags with *dnaA*, or plasmids with *repC*, and annotated with Prokka v1.14 [**Seemann, 2014**].

### Taxonomic order-wide genome tree construction and comparative genomics

To detect genes for photosynthesis among all cultured and uncultured *Rhizobiaceae* known to date, we screened the 17,766 genomes and MAGS of *Rhizobiales* currently available in the Genome Taxonomy Database (GTDB) v232 [**Parks *et al.,* 2025**]. First, the low-quality MAGS (checkM2 completeness >80%, contamination < 10%) were filtered out via BakRep v2 [**Fenske *et al.,* 2024**], which yielded 7527 of high-quality MAGS. A genome tree was then generated using GToTree v1.8.17 [**Lee, 2019**], linking Prodigal v2.6.3, HMMER3 v3.3.2, Muscle v5.1, TrimAl v1.4.rev15, and FastTree2 v2.1.11 as a pipeline to allow a multi-locus-sequence alignment (MLSA) of 117 single-copy protein-encoding genes that are conserved in *Alphaproteobacteria*, employing an approximate maximum-likelihood method to generate a phylogenetic tree.All 7527 genomes were further screened for nodulation (*nodA*), photosynthesis (*pufM*), and nitrogen fixation (*nifK*) genes via HMMER3 v3.4 search [**Eddy, 2011**], as shown in Table S4.

### Ancestral genome reconstruction and analysis of gene family dynamics in Rhizobiaceae

A family-specific phylogenomic tree was generated to place the newly sequenced genomes among recently differentiated *Rhizobiaceae* genera [**Kuzmanović *et al.,* 2022; diCenzo *et al.,* 2026**] (Table S5A). The 77 high-quality genomes contained 208 conserved single-copy protein-encoding genes (Table S5B) identified under the Codon-Tree strategy within PATRIC (BV-BRC) v3.57.40 [**Wattam *et al.,* 2017**] for subsequent RAxML analysis [**Stamatakis, 2014**]. Accessory and pangenomes of the genera, species, or unique clades of the family were identified via an upset plot using Proteinortho v6.3.6 [**Lechner *et al.,* 2011**] (Tables S5C, S5D). Gene family innovations, contractions, and ancestral genomes were then inferred across the species tree using the GLADE v1.0 pipeline [**Belcher and Kelly, 2026**]. Here, orthologous genes were clustered via OrthoFinder v3.1.5 [**Emms and Kelly, 2019**] and individual gene trees were simultaneously reconciled against the species tree topology. Every gain, duplication, and loss event could then be mapped per ortholog, with ancestral genomes reconstructed to each internal node, allowing for net genome expansion and contraction quantification across the evolutionary history of this bacterial family, while also tracking explicit gene development (Table S6).

Genus-specific genome trees focusing on distinguishing phototrophic MAGs related to *Peteryoungia* (Table S7) or *Hoeflea* (Table S8) were also built via PATRIC (BV-BRC) v3.57.40, to included all available environmental representatives sourced from MGnify [**Gurbich *et al.,* 2023**] in addition to those of GTDB, permitting strain-level analysis of gene presence/absence as compared to sample sources.

Heredity of the photosynthetic gene cluster (PGC) genes was further identified using circularized genomes in a progressive mauve alignment with automatic seed weight and minimum locally collinear block scoring, to reveal if each PGC had high synteny, suggesting vertical inheritance, or low synteny to infer horizontal gene transfer (HGT) [**Darling *et al.,* 2004**]. Photosynthesis genes as concatenated *bchXYZ*-*pufLMC* amino acid (AA) sequences were then compared to the evolutionary history of 16S rRNA genes via contrasting Maximum Likelihood trees employing optimal Le_Gascuel_2008 for AA [**Le and Gascuel, 2008**] and Tamura-Nei for nucleic acid modeling [**Tamura and Nei, 1993**], yielding highest log likelihood (−66359.09 and −16181.10) trees, respectively. Initial trees for the heuristic search were obtained automatically by applying Neighbor-Joining and BioNJ algorithms to a matrix of pairwise distances estimated using the JTT or Tamura-Nei model, respectively, and then selecting the topology with superior log likelihood value. A discrete Gamma distribution modeled evolutionary rate differences among sites (5 categories (+*G*, parameter = 0.4537 or 0.5597)), utilizing 1000 bootstrap iterations. A rate variation model allowed for some sites to be evolutionarily invariable ([+*I*], 27.18 or 8.15% sites, respectively). Both trees were drawn to scale, with branch lengths measured as the number of substitutions per site. Datasets involved 59× 1580 nucleotide and 2270 amino acid positions respectively. Evolutionary analyses were conducted in MEGA X [**Kumar *et al.,* 2018**].

### *Rhizobiaceae* evolutionary timeline calculations

A total of 181 genomes, including those generated for the *Hoeflea* and *Peteryoungia* strains were used for molecular dating based on phylogenetic reconstruction (Table S9A). Fossil records (Table S9B) anchored the ancestral age of specific *Alphaproteobacteria* and *Cyanobacteriota* linages [**Wang and Luo, 2021**], allowing for the extrapolation of *Rhizobiales* speciation events [**Wang *et al.,* 2020**]. Twenty universally conserved and validated bacterial single-copy core gene AA sequences with highest phylogenetic fidelity among bacteria were chosen [**Tian and Imanian, 2023**]. We employed Hidden Markov Model (HMM) profiles corresponding to target protein families downloaded from the InterPro database. These profiles were used to search against all genomes with HMMER v3.4, AA sequences were then aligned using MAFFT v7.490 [**Katoh *et al.,* 2019**], trimmed using TrimAl v1.5.0 [**Capella-Gutiérrez *et al.,* 2009**] and concatenated. For phylogenetic inference, each gene was treated as a separate partition for a best-fit substitution model using ModelTest-NG [**Darriba *et al.,* 2020**], restricting the search to the Le-Gascuel (LG), Whelan-Goldman (WAG), and Jones-Taylor-Thornton (JTT) approaches [**Jones *et al.,* 1992**]. The resulting scheme was used for maximum-likelihood tree reconstruction with RAxML-NG [**Kozlov *et al.,* 2019**] using the --all workflow and 500 bootstrap replicates. The phylogenetic tree was rerooted at midpoint via Newick Utilities [**Junier and Zdobnov, 2010**]. Estimates of divergence times were obtained via MCMCtree v4.10.9 [**Dos Reis and Yang, 2019**] of the PAML package [**Yang, 2007**], employing the concatenated AA alignment and a calibrated tree as input files. Branch lengths were modified via ancestral age calibrations (Table S9B), allowing for 2.5% posterior probability [**Wang *et al.,* 2020**]. The rate-prior parameter rgene_gamma was estimated, and an empirical LG AA substitution model was built with an independent-rates molecular clock. 10,000 iterations were used as burn-in, sampling every two iterations until 50,000 samples were collected. Phylogenies were visualized with iTOL v7.2.1 [**Letunic and Bork, 2024**].

### Biogeography and specific environments of different Rhizobiaceae lineages

The global presence and environmental conditions of sampling sites from where *Hoeflea* and *Peteryoungia* spp. Strains originated were compared to conditions under which *Rhizobium* was found during various surveys of 16S rRNA microbial communities as provided by the MicrobeAtlas v1.0 package [**Rodrigues *et al.,* 2017**]. Representative taxa which had sufficient sample coverage and metadata available were identified (Table S10A), listing general sample type categories (soil, aquatic, plant, animal) and geographical abundance maps via the MAP v3 visual interface. Moreover, the listed samples containing taxa of interest were screened, documenting the unique habitats each species of *Hoeflea* or *Peteryoungia* naturally resides at highest abundances (Table S10B).

## Results and Discussion

### Occurrence of anoxygenic phototrophy in *Rhizobiaceae*

Many of the classic plant-associated bacteria belong to the family *Rhizobiaceae* within the order *Rhizobiales*. The genus *Hoeflea* has recently been recognized as a member of this family [**Kuzmanović *et al.,* 2022; Naranjo-Robayo et al., 2026**], where *Hoeflea phototrophica* [**Biebl *et al.,* 2006**] and *H. olei* [**Rahul *et al.,* 2015**] are marine aerobic heterotrophs previously reported to contain bacteriochlorophyll (BChl) *a*, with the former also known to associate with dinoflagellates. Prior to such taxonomic rearrangements, ‘photosynthetic rhizobia’ referred only to *Bradyrhizobium* spp. [**Evans *et al.,* 1990; Fleischman and Kramer, 1998; Jaubert *et al.,* 2008; Avontuur et al., 2023; Ling *et al.,* 2024**]. However, the latter now belong to the *Nitrobacteraceae* and consequently share only a distant relationship to symbiotic *Rhizobiaceae*.

Notably, most species of *Hoeflea* associate to oxygenic phototrophic algae. Six alga-associated *Hoeflea* species are recognized, comprising *H. alexandrii*, *H. anabaenae*, *H. prorocentri*, *H. portis*, *H. ulvae*, and *H. algicola* (Table S1). Of these, *H. anabaenae* WH2K exclusively attaches to *Anabaena variabilis* SSM-00 heterocysts [**Stevenson and Waterbury, 2006**] (Fig. 1A,B) and is supplied with fixed carbon and nitrogen by its cyanobacterial host [**Behrens *et al.,* 2008**], while the newly isolated *H. alexandrii* DSM 118613 is found to bind *Dunaliella bicculata* SAG 19-4 in coculture (Fig. 1C,D). When characterizing novel strains, the capability of aerobic anoxygenic phototrophy can be overlooked if growth conditions suppress the expression of photosynthesis genes, leaving the cells unpigmented [**Hughes *et al.,* 2018**]. Excess vitamin and trace metals, low organic carbon [**Biebl and Wagner-Döbler 2006; Kuzyk *et al.,* 2023**], or decreased salt concentrations [**Biebl *et al.,* 2006**] have been shown to stimulate BChl *a* production. Indeed, we observed that axenic *H. anabaenae* WH2K developed a fuchsia-pink colour when grown in PY+ medium supplemented with trace elements and vitamins (Table S2), in clear contrast to the cultures with standard PY lacking traces or copiotrophic 2216 marine broth (Fig. 2A). We therefore systematically investigated other AAP in *Rhizobiaceae*, especially those associated with cyanobacteria, algae, or plants.

**Fig. 1.**
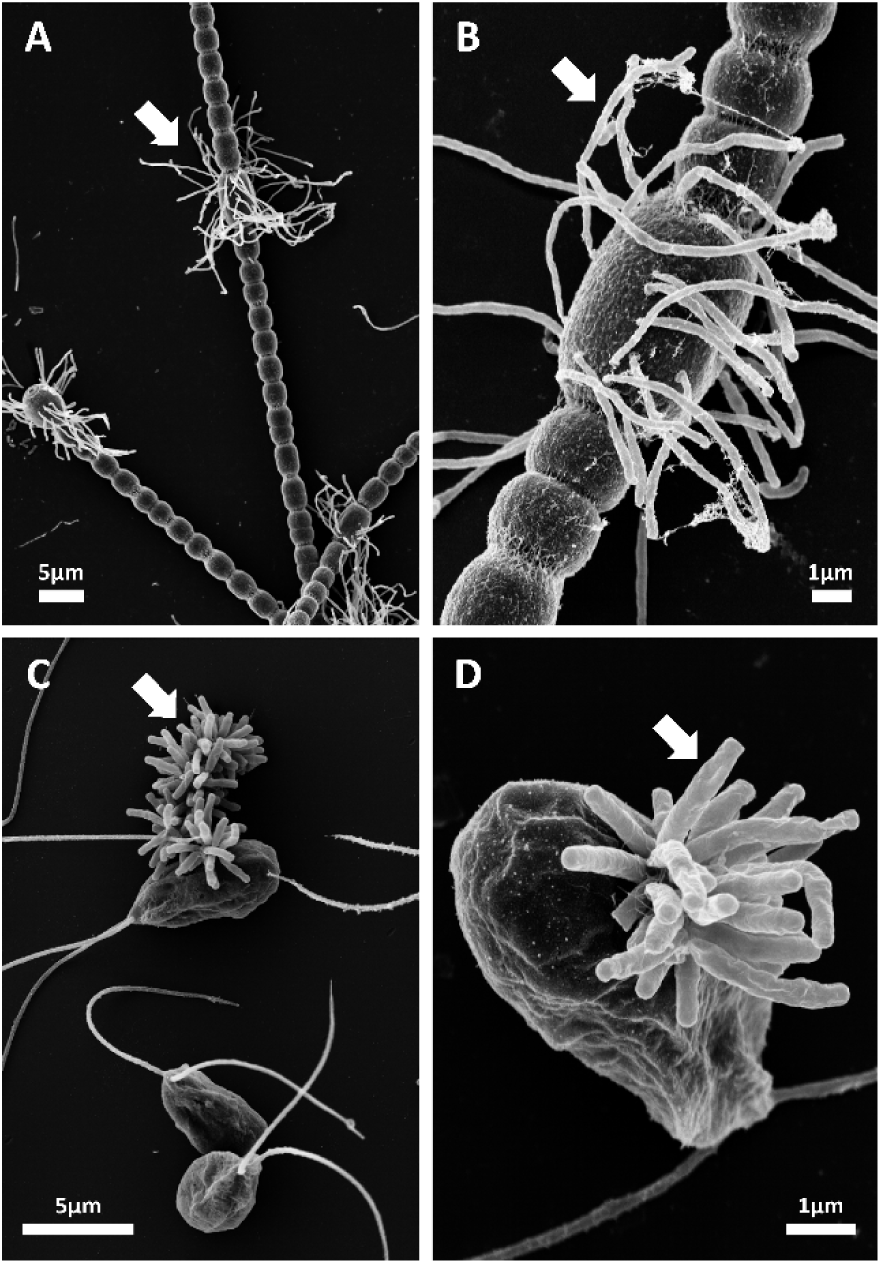
*Hoeflea* spp. bind microbial oxygenic phototrophic cyanobacteria or algae as epibionts. **(A,B)** Scanning electron micrographs of *Hoeflea anabaenae* WH2K cells selectively attached to nearly all heterocysts of *Anabaena variabilis* SSM-00, while **(C,D)** *Hoeflea* sp. DSM 118613 bind the surface of several *Dunaliella bicculata* SAG 19-4 in mixed culture. Epibiont *Hoeflea* spp. cells, white arrows.

**Fig. 2.**
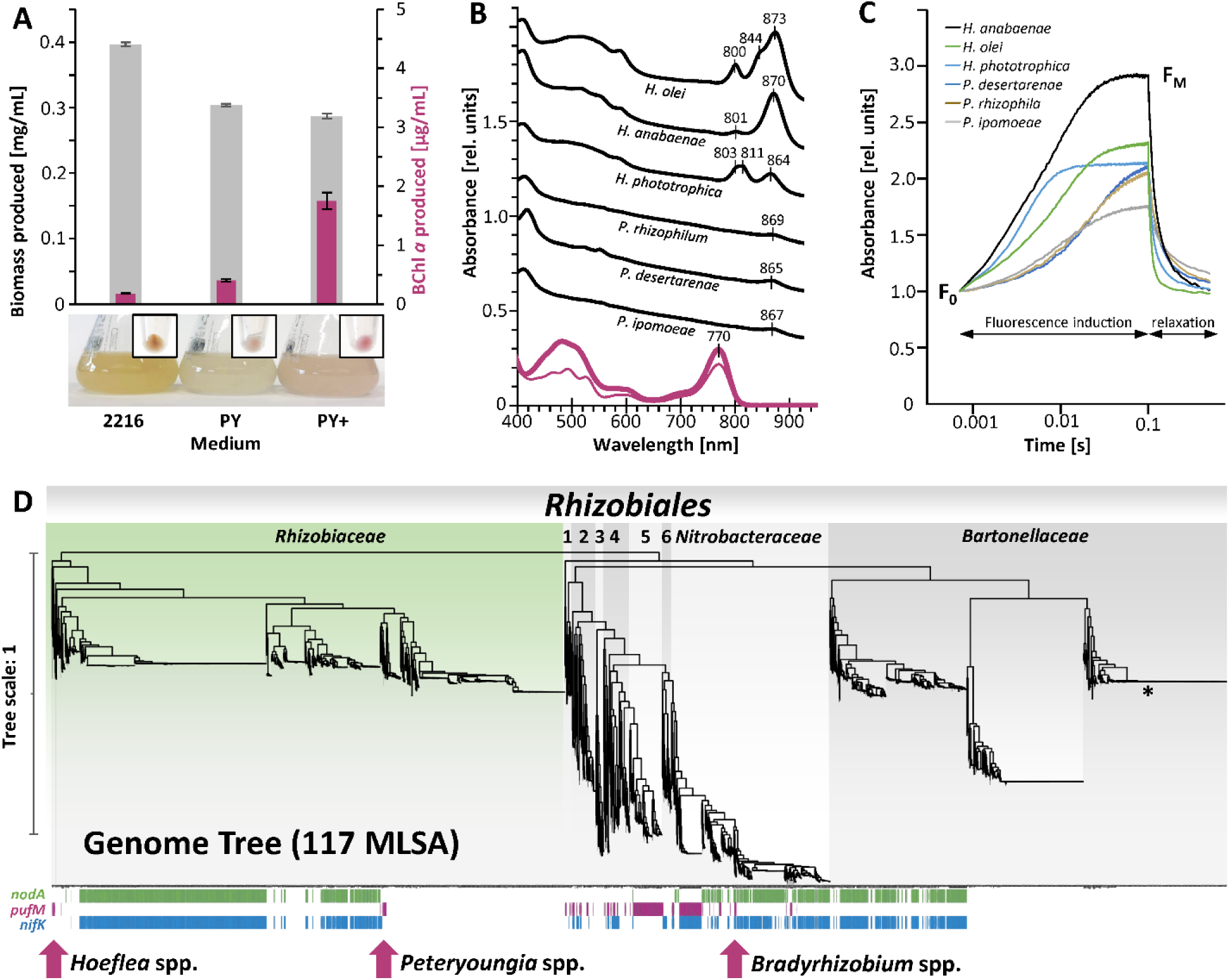
Photosynthetic complexes and pigments discovered in diverse *Rhizobiaceae* include *Hoeflea* and *Peteryoungia*. **(A)** *H. anabaenae* exhibits a genus-wide trend of pigmentation only under nutrient poor conditions (media PY and PY+), stimulated by presence of trace metals and vitamins (medium PY+), otherwise inhibited under standard rich organic Marine 2216 medium. **(B)** Absorbance spectra of phototrophic *Hoeflea* and *Peteryoungia* representatives compare whole-cells and pigment extracts revealing light harvesting and reaction centers that absorb near-infrared, where *H. phototrophica* has an unusual LHII peak at 811 nm, and *H. olei* has a LHII shoulder at 844 nm. All strains have BChl *a* maxima in extracts at 769 nm, with spectra at 400-600 nm show *H. olei* and *H. phototrophica* to have spheroidenenone carotenoids (bold pink trace) while all others contain spirilloxanthin (thin pink trace). **(C)** Light-induction fluorescence kinetic assays revealed photosynthetic activity of all tested *Hoeflea* and *Peteryoungia* spp. **(D)** Genome tree of all 7527 high quality *Rhizobiales* representatives sourced from GTDB via BacRep differente taxonomic families, and presence of nodulation (*nodA*), photosynthesis (*pufM*), or nitrogen fixation (*nifK*) genes. Only *Hoeflea* and *Peteryoungia* relatives contain photosynthesis genes in *Rhizobiaceae*, while classically described *Bradyrhizobium* relate to *Nitrobacteraceae*. Notable family compositons: 1, *Ahrensiaceae*; 2, *Afifellaceae* & *Stappiaceae*; 3, *Devosiaceae*; 4, *Hyphomicrobiaceae*, *Beijerinckiaceae* & *Boseaceae*; 5, *Methylobacteriaceae*; 6, *Xanthobacteraceae*. * 2611× *Bartonella melitensis* strains collapsed to allow visualization of other families/species, since no *B. melitensis* contain nodulation, photosynthesis, or nitrogen fixation genes.

Indeed, many of the 7527 high-quality *Rhizobiales* genomes available in GTDB were shown to contain photosynthesis-related genes, as compared to nodulation or nitrogen fixation genes (Table S4), mapping to various lineages including *Rhizobiaceae* (Fig. 2D). While several draft genomes of *Hoeflea* spp. putatively contained photosynthesis genes, multiple relatives of distantly related *Peteryoungia* were also positive (Table S1, S4). For a deeper phylogenomic analyses, we newly sequenced 12 genomes of *Hoeflea* and *Peteryoungia* type strains, which yielded high quality, circularized chromosomes and up to two plasmids per genome (Table S3, Fig. S1). Photosynthesis genes were indeed detected in *H. phototrophica, H. olei, H. alexandrii*, *H. anabaenae*, *P. ipomoeae*, *P. rosettiformans*, *P. glycinendophyticum*, *P. desertarenae*, *P. albertimagni*, *P. aggregata*, and *P. rhizophila,* with the latter nine identified for the first time by our study (Table S1). In fact, no single *Peteryoungia* species had previously been recogenized as an AAP, with all members originally described as nonpigmented heterotrophs. The genes encoding photosynthetic reaction center and antenna proteins as well as pigment synthesis pathways were always located on chromosomes and were organized in Photosynthetic Gene Clusters (PGCs) that are characteristic for the majority of anoxygenic photosynthetic bacteria (Figs. S1, S2). No PGC was found on plasmids as seen in other *Alphaproteobacteria* families [**Zheng *et al.,* 2011**; **Liu *et al.,* 2019**].

Each suspected AAP was tested by biochemical and physiological methods in pure culture to confirm the expression of this trait. In parallel, all available information on these isolates concerning their potential association with other organisms was gathered from literature and databases (Table S1). *In vivo* spectroscopy indicated the formation of intact photosynthetic complexes in every tested *Peteryoungia* spp., as well as in the four *Hoeflea* species *H. alexandrii, H. anabaenae H. olei, and H. phototrophica* when grown until stationary phase on agarized PY+ medium (Fig. 2B, S3). We found that excess trace elements stimulated BChl *a* production more effectively than vitamin addition in all tested strains (Fig. S4A), as did reduced salt concentrations of 1-2% (Fig. S4B), suggesting niche quorum sensing. While most strains preferred lower concentrations of organic carbon compounds for their pigment production as seen previously [**Kuzyk *et al.,* 2023**], only *H. olei* uniquely expressed BChl *a* constitutively (Fig. S4C). Each strain required aerobic conditions and anaerobic growth was not detected.

HPLC analysis of pigment extracts detected BChl *a* esterified with phytyl side chains (BChl *a*_p_ (Fig. S5), in all *Hoeflea* spp. and *Peteryoungia* spp. except *H. anabaenae*. The latter contained BChl *a* esterified with geranylgeranyl, dihydrogeranylgeranyl, tetrahydrogeranylgeranyl besides phytyl. The carotenoids synthesized by *Peteryoungia* strains were those of the spirilloxanthin pathway, the deep branching *H. anaebaenae* contained spirilloxanthin and an unknown ketocarotenoid with an absorption maximum 493 nm (Fig. S2B, S5), whereas *H. phototrophica* and *H. olei* produced hydroxyspheroidenone, spheroidenone, and an unknown minor carotenoid. The genes of each PGC matched carotenoids observed, as all *Peteryoungia* spp. had *crtIBCDEF* which are indicative of spirilloxanthin carotenoids [**Sandmann, 2024**] (Fig. S2A). In contrast, *Hoeflea phototrophica* and *H. olei* contained the marker gene *crtA* encoding spheroidene monooxygenase [**Sandmann, 2024**] which is consistent with the presence of spheriodenone in these species. Although the partially inverted PGC of *H. anabaenae* matched some structural aspects of those found in distantly related *Bradyrhizobia* (Fig. S2A), its lack of *crtA* was in-line with the predominance of detected spirilloxanthin, while the distal position of *crtIB* separated with two unknown genes may explain its unusual minor carotenoids. The photosynthetic gene contents also mirrored the photosynthetic complexes inferred from *in vivo* spectroscopy. In addition to a standard reaction center (RC) and light harvesting complex (LH) I that typically feature absorption maxima at ∼802 and 867 nm, *H. phototrophica* and *H. olei* presented respective absorption maxima or shoulders at 811 and 844 nm indicating LHII (Fig. 2B, S3), matching the presence of additional *pucABC* genes in both strains (Fig. S2A). Every pigmented *Hoeflea* and *Peteryoungia* spp. was further shown to utilize their photosynthetic machinery, as fluorescence transients confirmed each RC to be photochemically active (Fig. 2C). Based on the combined genomic physiological, spectroscopic and photochemical analyses, it is evident these diverse *Rhizobiacae* possess phototrophic capabilities as AAP.

### Evolution of aerobic anoxygenic photosynthesis in *Rhizobiaceae*

Our unexpected discovery of numerous previously unknown AAPs in *Rhizobiaceae* prompted an investigation into the evolutionary origin and persistence of photosynthesis during the radiation of this typically plant-associated family. Many *Pseudomonadota* species, particularly in *Alphaproteobacteria*, contain BChl *a* photosystems. Congruent phylogenetic patterns of core housekeeping and photosynthesis genes across the class have revealed a shared ancestral origin of this metabolic trait, which was subsequently lost in multiple lineages [**Imhoff, 2017; Tinguely *et al.,* 2023; Imhoff and Kyndt, 2026**]. Moreover, the successful and complete acquisition of about 50 photosynthesis genes >45 kbp by HGT is considered a rare event. Yet, phylogenomic evidence suggests that HGT has occurred within the *Alphaproteobacteria* among *Rhodobacteraceae* spp., infrequently between members of the classes *Alpha*-, *Beta*-, and *Gammaproteobacteria* [**Brinkmann *et al.,* 2018**], and rarely across the borders of bacterial phyla from *Pseudomonadota* to *Gemmatimonas phototrophica* of the phylum *Gemmatimonadota* [**Zeng *et al.,* 2014**]. Such HGT events have been linked to the presence of complete PGCs on plasmids, and potential transformations [**Brinkmann *et al.,* 2018**]. Many *Rhizobiaceae* also contain plasmids which likely mediated the occasional HGT of nodulation factors or *vir* genes for mutualistic and antagonistic interactions, respectively [**Wang *et al.,* 2019; Wardell *et al.,* 2021**]. Similarly, the PGC of closely related *Bradyrhizobium* species were suspected to be acquired by HGT [**Giraud *et al.,* 2000; Avontuur *et al.,* 2023**]. For the first time, our complete genomes revealed plasmids in three *Hoeflea* and four *Peteryoungia* type strains (Table S3), but each PGC was found chromosome bound (Fig. S1).

To reconstruct an evolutionary history of anoxygenic photosynthesis in *Rhizobiaceae* we incorporated the new genomic data into a GLADE analysis utilizing orthologous gene identification [**Belcher and Kelly, 2026**], tracing all gene gain, loss, and duplication events to assemble predicted ancestral genomes (Table S6, Fig. 3). By following *pufM* as a representative essential gene of anoxygenic photosynthesis, we find the last common ancestor (LCA) at the root had photosynthesis genes (Table S6B), with multiple lineages consecutively losing this trait (Fig. 3; Fig. S6). A similar trend was observed for nitrogen fixation genes with *nifK* depicted, where these genes are also predicted as an ancestral trait which suffered massive gene loss, as reported elsewhere [**Yang *et al.,* 2020**; **Sobol *et al.,* 2026**]. In comparison, nodulation genes such as *nodA* instead appear after the first recognized split of the *Rhizobiales*, but then were vertically inherited, with multiple *Rhizobiaceae* lineages also losing this trait (Fig. 3). This finding also matches expected nodulation inheritance, where *nod* factors may have developed in *Rhizobiaceae* around 51 Mya following plant evolution, with consecutive HGT into *Nitrobacteraceae* (*Bradyrhizobium*) and *Paraburkolderia* as separate events [**Rahimlou *et al.,* 2021**].

**Fig. 3.**
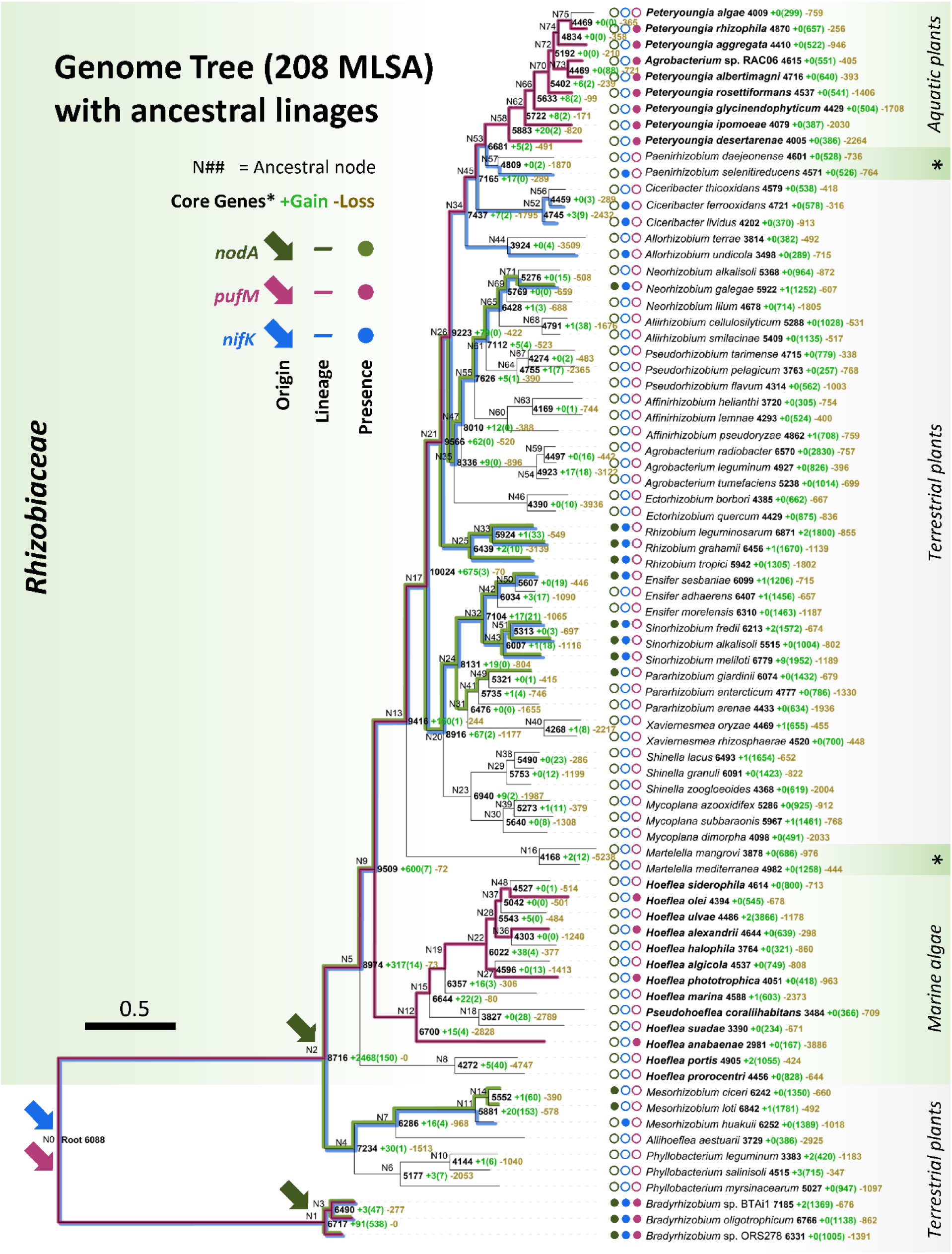
Phylogenomic relationships with ancestral gene gains and losses within the *Rhizobiaceae*. Ancestral state analysis superimposed on a genome tree built with 208 core genes of 77 genomes covering all recently proposed genera of the *Rhizobiaceae* [**Kuzmanovic *et al.,* 2022; Naranjo-Robayo et al., 2026**], utilizing *Afifellaceae, Phyllobacteriaceae* and *Nitrobacteraceae* members as outgroups (Tables S5A,B). Evolutionary history was retraced for every orthologous gene among all genomes via GLADE analysis, predicting gene gain and duplication (bright green), and loss (dark yellow) events to reveal gene origin, lineage, and presence among each type species and ancestral nodes. Nodulation (dark green), photosynthesis (maroon), and nitrogen fixation (blue) were followed via *nodA*, *pufM*, and *nifK*, respectively. The inferred oxygenic phototrophic hosts are listed on the right as terrestrial plants, aquatic plants, marine algae, or unknown aquatic (asterix).

As the evolutionary history reconstruction suggested that the LCA of *Rhizobiaceae* was a phototroph, we tested this hypothesis through comparison of conserved 16S rRNA genes to concatenated AA sequences of *bchXYZ-pufLMC* from all photosynthetic representatives (Fig. 4A). Phylogenetic congruence was observed via similar branching patterns, with family members clustering separately from outgroups, and *Peteryoungia* vs. *Hoeflea* spp. differentiating into similar clade structures in both trees (Fig. 4A, S7A). The conclusion of a common ancestral origin of photosynthesis genes in *Rhizobiaceae* was further supported by consistent GC% values in the neighborhood of the PGC on the chromosomes ruling out recent recombination events (Fig. S1), and by the high levels of genome synteny including PGC location (Fig. 4B, S7B). *P. algae* SSM4.3 is the sole *Peteryoungia* strain lacking a PGC, where a secondary loss can be inferred from the synteny of up and down stream genes as compared to closely related *P. aggregata* and *P. rhizophila* (Fig. 4C). A similar loss of PGC is also evident for *Hoeflea* spp., where flanking genes remaining syntenous with those of its immediate relatives (Fig. S7C). The observation of *H. anabaenae* WH2K as the deepest branch of *Hoeflea* clade is in-line with the detection of common denominator spirilloxanthin carotenoids and BChl *a* homologs with different esterifying isoprenoid alcohols in this lineage (see above). While the genome of this strain showed an inverted section of its PGC mirroring *Bradyrhizobium* spp. (Fig. S2), our analysis indicates this to have occurred independently across phylogenetic lineages. Based on all our phylogenomic analyses, the currently known AAP *Rhizobiaceae* likely evolved from a common phototrophic ancestor rather than acquiring photosynthesis genes independently and through multiple recent HGT events.

**Fig. 4.**
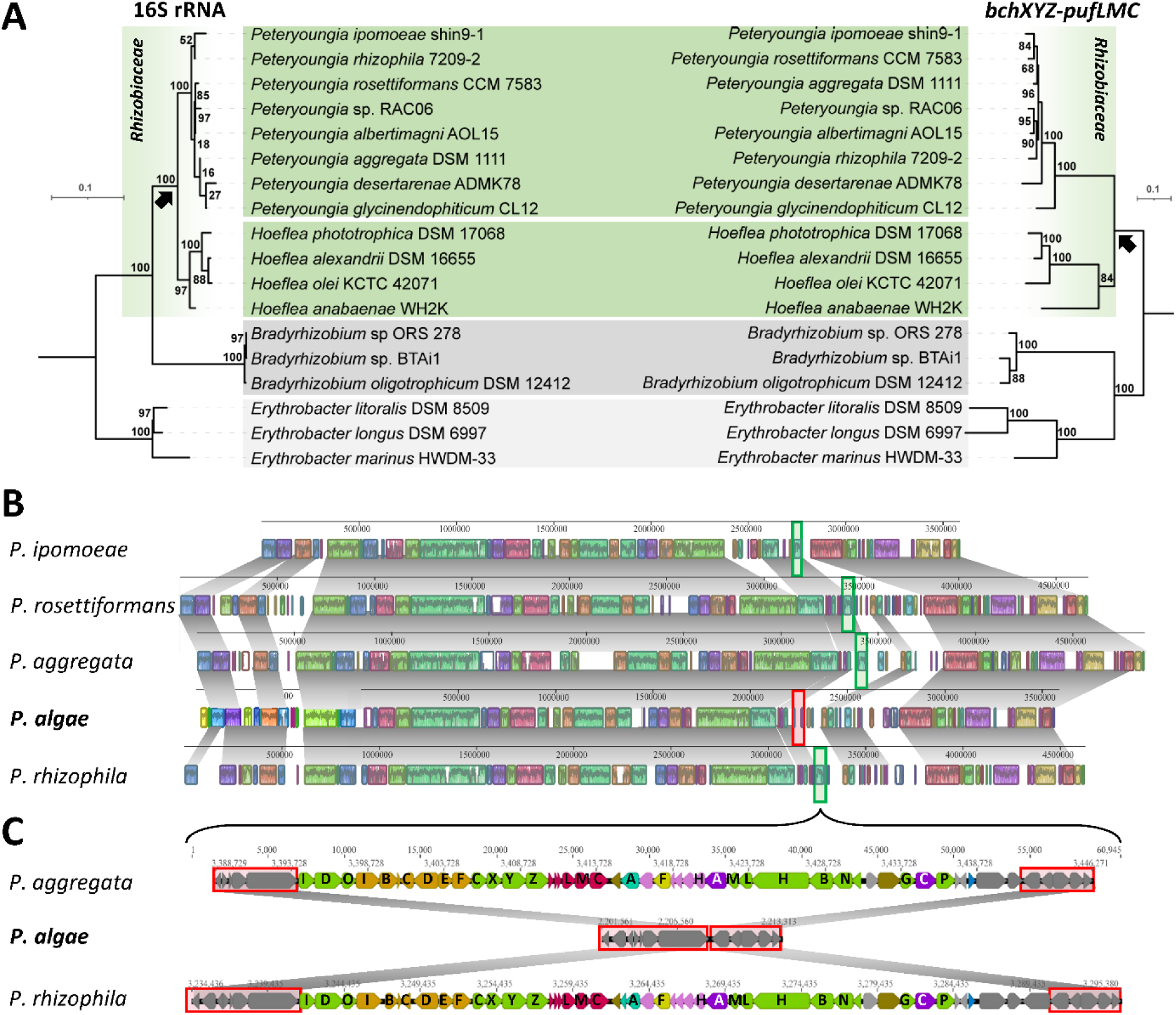
Vertical inheritance and progressive gene loss of phototrophic *Rhizobiaceae* shown by genome synteny and gene homology patterns. **(A)** Phylogenetic evolution comparison of 16S rRNA gene homology vs. *bchXZY-pufABC* tandem genes of *Rhizobiaceae* members compared to *Bradyrhizobium* and *Erythrobacter* as outgroups. Similar branching patterns suggests all *Rhizobiaceae* members (black arrow) to have contained photosynthesis for a long evolutionary period as similar genetic drift occurs as for 16S rRNA genes. **(B)** High collinearity exists for all *Peteryoungia* spp. as mapped with MAUVE alignment, showing identical locations of PGC (green box) in genome as compared to other genetic elements. **(C)** Comparing related phototrophic species *P. aggregata* and *P. rhizophila* with *P. algae* genome reveals the lack of PGC, but retains flanking genes (red boxes).

### Co-evolution of *Rhizobiaceae* with cyanobacteria, algae and plants

Next, we identified the likely drivers of massive gene loss, and the possible selective advantages that promoted few members of *Hoeflea* and *Peteryoungia* to maintain this feature. Multiple independent losses of ancestral phototrophy in AAP have been proposed among other *Pseudomonadota* families [**Koblížek *et al.,* 2013; Kasalický *et al.,* 2017**], attributed to the significant energetic burden of AAP photosynthetic apparatus biosynthesis [**Kirchman and Hanson 2013**; **Tinguely *et al.,* 2023**], that would be counter selected and lost upon adaptation to a new mostly or permanently dark niche.

Examining the habitats of diverse cultured *Rhizobiaceae* revealed a clear ecological pattern of phototrophic members predominantly found in illuminated aquatic habitats, whereas obligate chemotrophs were found in terrestrial soils (Fig. 3). Moreover, all species are strongly associated to specific oxygenic phototrophs in various forms of symbiosis, with terrestrial *Rhizobiaceae* often developing root nodules, *Peteryoungia* strains were frequently observed on submerged roots of aquatic plants (Fig. S8), whereas *Hoeflea* spp. occupied the phycosphere directly bound to aquatic algae (Fig. S9). Analysis of culture-independent microbial community studies on a global scale supported both generalized habitat and specific symbiont associations, as shown by the relative number of samples identified in Microbe-Atlas sorted by reference sequence (Fig. S10). It is evident that *Rhizobium leguminosarum* is predominant in soil habitats (4296 samples), particularly fields (411), whereas *P. rosettiformans* is often connected to plants (1420), *H. phototrophica* is found in aquatic habitats (781), and *H. anabaenae* as the rarest group has a mixture of general sample types (Table S10a). The highest abundant samples provided additional clarity to each symbiotic association (Table S10b), with *R. leguminosarum* affiliated with legumes (>29000 ppm), *P. rosettiformans* often found on submerged plant roots (>1700 ppm), *H. phototrophica* with saltwater algae (>2000 ppm), and *H. anabaenae* predominantly linked to cyanobacteria samples (>1000 ppm). Together, type-species isolation sources, metagenome locations, and global 16S rRNA gene libraries suggest each genera is indeed associated to different, yet specific, oxygenic phototrophs.

To better understand how the development of symbiotic relationships may have impacted the conservation of phototrophy and differentiation of *Peteryoungia* from *Hoeflea*, we built a time-scale evolutionary phylogenetic tree to follow *Rhizobiaceae* development as compared to origins of oxygenic phototrophs (Fig. 5). Although rapid changes via HGT and a lack of direct classical fossil evidence for multiple lineages can make it difficult to estimate bacterial evolutionary timescales [**Liuca *et al.,* 2018**; **Wang and Luo, 2021**], the ancestral age of *Rhizobiales* can be estimated based on the diversification on hosts/ symbiovars [**Chriki-Adeeb and Chriki 2016**; **Wang *et al.,* 2020**; **Wang *et al.,* 2021**], supported by evidence of deep branching *Cyanobacteriota* evolution [**Wang *et al.,* 2020**; **Dos Reis and Yang, 2019**]. Calibrated with fossil records (Table S9), the ancestral age of each taxon was explored as it related to the evolution of legumes, land plants, or eukaryotic algae (Fig. 5). In this way, *Alphaproteobacteria* are predicted to have diverged 1.934 Bya while *Rhizobiales* evolved 1.517 Bya ago. Of *Rhizobiaceae*, the *Hoeflea* LCA branch 707.7 Mya, which dates before the proliferation of eukaryotic algae 610-660 Mya [**Brocks *et al.,* 2017**], while *Hoeflea* associated with algae diverge 550.3 Mya. In comparison, *Peteryoungia* occur 270.1 Mya, after adaptation of land plants 450-500 Mya [**Morris *et al.,* 2018**], but prior to the development of legumes 65-110 Mya [**Barba-Montoya *et al.,* 2018**], when several nodulating lineages appear across *Rhizobiaceae* (Fig. 5, black dots). The deepest branching *Rhizobiaceae* phototroph is *H. anabaenae* 435 (Fig. 5), a species capable of direct adhesion to prokaryotic cyanobacteria [**Stevenson and Waterbury, 2006; Stevenson *et al.,* 2011**]. This lineage may be a remnant, or even a prokaryotic “living fossil” of the earliest form of symbiosis between *Rhizobiaceae* and oxygenic phototrophs, maintaining associations to simpler oxygenic phototrophs as recently reported for ‘*Ca.* Tectiglobus spp.’, a deep-branching member of the *Rhizobiales* [**Tschitschko *et al.,* 2024**].

**Fig. 5.**
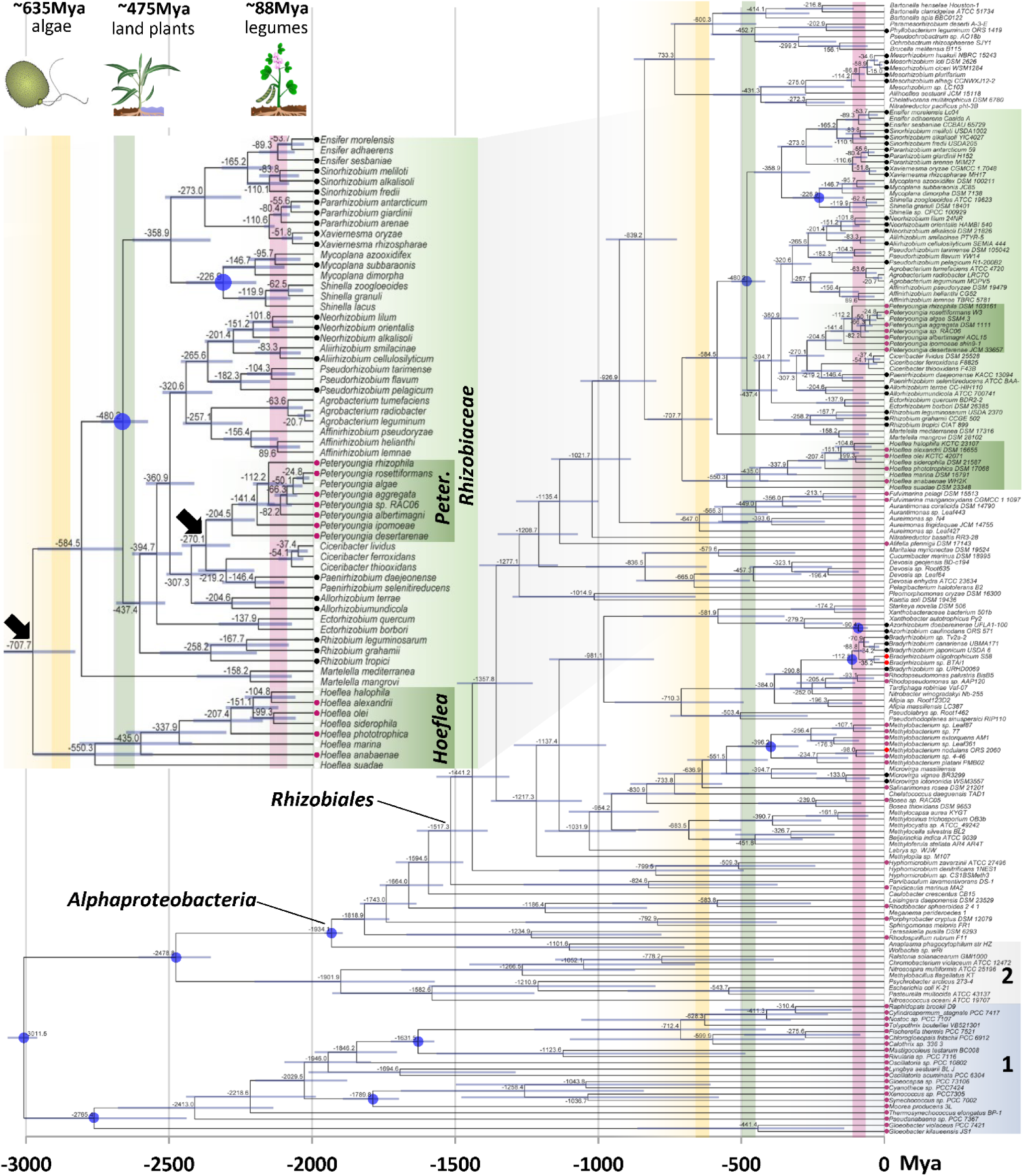
Time-scale evolutionary history of the *Rhizobiaceae* among *Alphaproteobacteria*, compared to predicted origins of oxygenic phototrophs. Divergence times were estimated using MCMCTree on the species phylogeny shown in Fig. 3 with additional members of *Pseudomonadota* and *Cyanobacteria*, totaling the 181 representative species. Nodes marked with a blue circle represent the fossil-record calibration points used to calculate ancestral linages (Table S9B), with delineation marked at each node with deviation as 95% HPD interval of posterior dates (blue bars). Vertical bars in yellow, green and red indicate the origins of algae, land plants, and legumes, respectively. The inset panel shows a zoomed-in view of the evolutionary timeline of *Rhizobiaceae*, where the hosts are also represented next to the tips of the phylogeny. *Cyanobacteriota*, 1; *Gammaproteobacteria*, 2; Phototrophs, maroon dots; nodulating bacteria, black dots; nodulating phototrophs, red dots; arrows, nodes of interest.

Combined, these results indicate bacterial speciation among this family occurred stepwise following eukaryotic oxygenic phototroph evolution. Moreover, this coevolution appears as a causative link to the evolutionary patterns that shape the presence or absence of photosynthesis in the developing *Rhizobiaceae* members, where only the taxa that stem from ancient epibiotic associations to aquatic plants or algae maintained this functionality. Additional support comes from *Methylobacteria*, another clade diverging during land plant radiation 551.5 Mya (Fig. 5). Leaf associated strains in this group predominantly harbor PGCs as documented elsewhere [**Zervas *et al.,* 2019**], a symbiotic location conductive to the maintenance of photosystems [**Grossi *et al.,* 2025**].

### Preadaptation of *Rhizobiaceae* for interactions with oxygenic phototrophs

By comparing the gene complement of both time-scale and ancestral gene gain/loss evolutionary trees we sought to elucidate the genetic basis of symbiosis between newly sequenced photosynthetic rhizobia and oxygenic phototrophic cyanobacteria, algae or plants. While genes for photosynthesis, nitrogen fixation, and flagellated motility are all found at the root LCA of *Rhizobiales* (Fig. 6), and several downstream clades loose one or all three traits (such as *Martellaceae*) (Fig. 3, Table 6B), the 839.2 Mya LCA of *Rhizobiaceae* and *Bartonellaceae* acquired a large number of genes (Fig. 3, Table 6C), including several for cell-signaling nodulation or virulence factors, vitamin production, and metal acquisition (Table S4b, S5c). While multiple pre-adaptations likely occurred at these ancestral nodes to permit associations with oxygenic phototrophs, nod-factors are of particular interest as they initiate the interaction of bacteria with the oxygenic phototrophic host, whereas downstream communication develops endosymbiosis. Thus, surface adhering epibionts (Fig. 1) could hypothetically utilize such nod-signal exchanges. Indeed, *nodMNG* were found in all tested *Rhizobiales* (Table S5A), where every *Hoeflea* has specifically maintained *nodEF,* which can yield nod-factors with altered fatty acyl chains [**Fisher and Long 1992; Patra and Mandal, 2022**]. The frequent presence of *nodT,* suggests some preadaptation among early branching members to produce the precursor sugar subunits of nod-factors [**Fisher and Long 1992; Batstone *et al.,* 2022**]. As atypical nod-factors have been found in *Bradyrhizobium* [**Bek *et al.,* 2010**], this may indicate that *Hoeflea* have unique signalling molecules to communicate with *Anabaena* or microalgae. In addition, virulence and adherence factors were ubiquitous across *Rhizobiaceae,* with *virA-B4-B11* genes known from *Agrobacterium* as type IV sectrion components [**Manksy *et al.,* 2022**] and adhesins also found (Table S5A). In support of commensal symbiosis, most *Rhizobiaceae* including the phototrophic *Hoeflea* and *Peteryoungia* have genes for niacin (B_3_), biotin (B_7_) and cobalamin (B_12_) generation (Table S5A), which are known to support the oxygenic phototrophic growth in other algal symbioses [**Cooper *et al.,* 2019**]. Moreover, all have iron efflux pump *fieF*, and some contain siderophore transport systems, indicating support in acquiring trace elements. Taken together, deep-branching *Rhizobiaceae* appear to possess several pre-adaptations for mutualistic associations with oxygenic phototrophs, wherein classical nod-factors may exert functions beyond plant root signaling to potentially encompass interactions with microalgae and cyanobacteria.

**Fig. 6.**
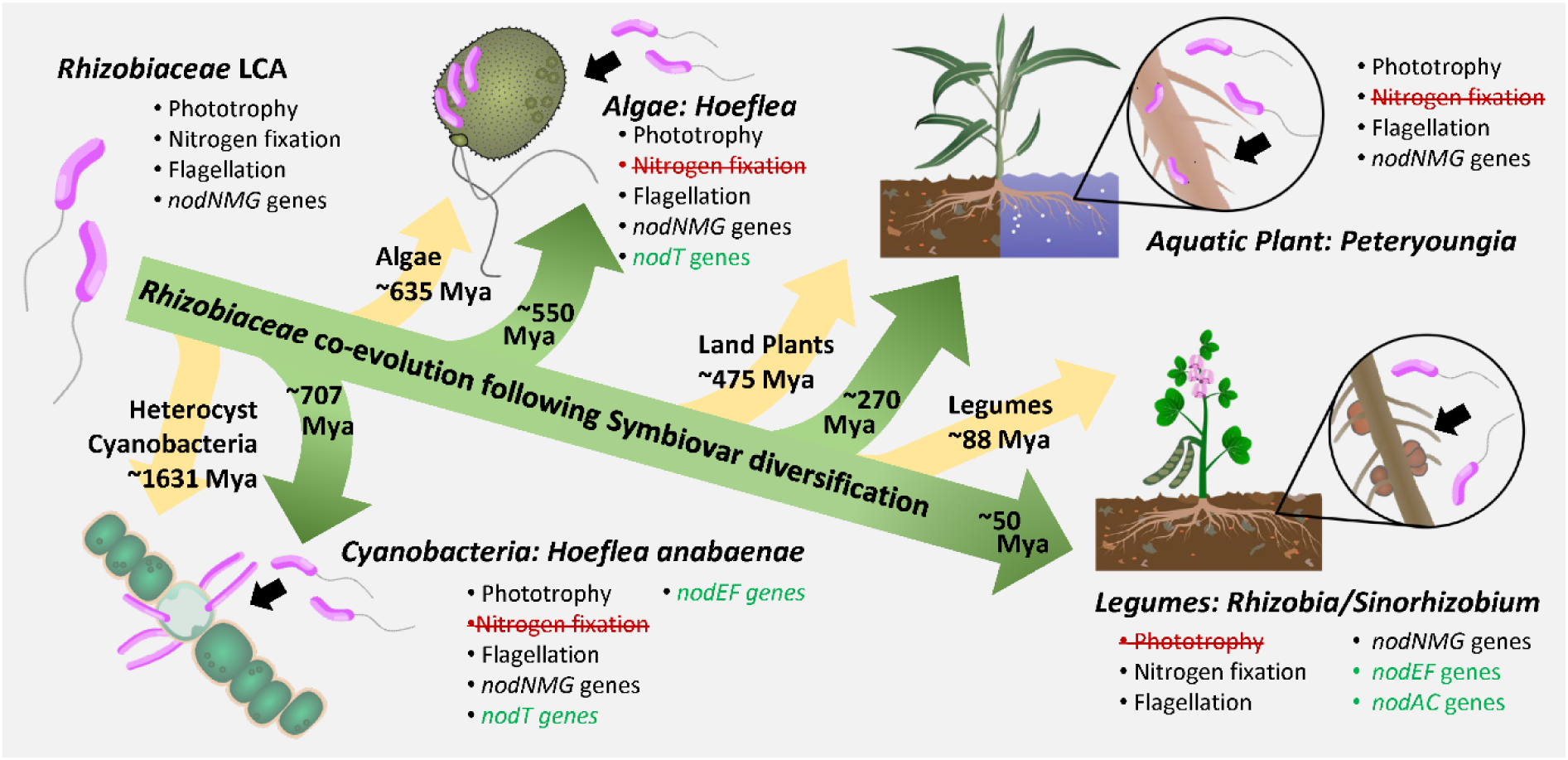
The coevolution of phototrophic Rhizobia as partners to specific oxygenic phototrophs. A schematic representation of the evolutionary trajectory of *Rhizobaceae* lineages following oxygenic phototroph evolution, as found by the diversification of the family (Fig. 2D), ancestral gene gains and losses (Fig. 3), vertical inhertiance and progressive gene loss of phototrophy (Fig. 4), and time-scale evolutionary histroy as compared to various oxygenic phototrophs (Fig. 5). Together, specific clades appear after the partnering host in the evolutioanry time-line, with specific gene content also being gained or lost dependent on the niche. It is proposed that *H. anabaenae* consortia with *Anabaena variabilis* [**Stevenson and Waterbury, 2006**], *H. phototrophica* isolated from *Prorocentrum spp.* [**Biebl *et al.,* 2006**], and *Peteryoungia* associated to aquatic macroalgae and plants such as *Ipomoea aquatica* [**Rahi *et al.,* 2021**], represent potential “living-fossils” from various points in the timeline.

## Conclusions

AAPs are facultative photoheterotrophs typically recognized as an ecologically relevant group of free-living bacteria found in upper layers of aquatic environments, the phyllosphere, soil crusts and also thermal springs [**Yurkov and Beatty, 1998; Koblížek, 2015; Tinguely *et al.,* 2023**]. In such environments, phototrophy has been shown to increase the fitness of AAPs when organic growth substrates become scarce [**Kirchman and Hanson 2013; Tinguely *et al.,* 2023; Kuzyk *et al.,* 2023**]. Many *Alphaproteobacteria* also associate with different cyanobacteria and algae where they provide vitamins or detoxify H_2_O_2_ in exchange for organic substrates [**Wagner-Döbler *et al.,* 2010; Kim *et al.,* 2019; Mironov *et al.,* 2019**]. As long as *Bradyrhizobium* spp. represented the only known plant symbionts capable of anoxygenic photosynthesis, the evolutionary origin and significance of this metabolic trait in *Rhizobiaceae* remained unclear.

Discovering descendants of phototrophic *Rhizobiaceae* associate with early branching lineages of oxygenic phototrophs, raise the question of additional ancestral traits that might have enabled these bacteria to associate with their hosts. An exchange of metabolites and growth factors appears relevant in the particularly well studied case of *Dinoroseobacter shibae*, with vitamins B_1_ and B_12_ produced by the APP in exchange of organic carbon substrates, niacin (B_3_), biotin (B_7_) and folate (B_9_) precursors excreted by the algal host [**Wagner-Döbler et al. 2010; Cooper *et al.,* 2019**], where the *Rhizobiaceae* epibionts reported here appear capable of producing niacin and biotin. Thus, while vitamin exchange may occur, it is presently unclear what adaptive value these AAP have in the symbiotic context.

However, as facultative photoheterotrophs, these AAP likely allow for a higher fraction of organic carbon substrates to be diverted towards anabolism, as chemiosmotic energy conservation is supplemented by light through the bacterial photosystem [**Biebl and Wagner-Döbler 2006; Holert et al. 2011**]. This metabolism has been found to be of particular advantage during diurnal light-dark cycles [**Soora and Cypionka 2013; Tinguely *et al.,* 2023**], and all *Rhizobiaceae* were further found to have circadian rhythm regulator *kaiC*. Thus, a key selective advantage of AAP symbionts could be the efficient utilization of host-derived organic carbon for anabolism, and less organic carbon burden as an obligate epibiont when compared to strict heterotrophic symbionts. Phototrophy can thereby be considered relevant not only for supplementing the energy budget of free-living *Alphaproteobacteria*, but based on our study also of selective advantage in the epibiotic symbiosis of bacteria with oxygenic phototrophs across the different domains and kingdoms of life.

## Acknowledgements

This research was funded by a grant of German Research Foundation (DFG OV 20/10-2). MKS and MK were supported by the project Photomachines (CZ.02.01.01/00/22_008/0004624) financed by the OP JAK program of the Czech Ministry of Education.

Sequencing via PacBio Sequell *IIe*, Oxford Nanopore, and Illumina NextSeq2000 was performed by the Sequencing Core Facility of the Services Department at the Leibniz Institute DSMZ. Anika Methner and Franziska Burkart are sincerely thanked for their assistance in preparing physiological assays; Victoria Ringel supported the deposition of *Hoeflea* sp. DSM 118613. We thank Ina Brentrop for EM sample preparation. Strains relating to *Peteryoungia* originating from India were accessed with permission from the National Biodiversity Authority, an autonomous and statutory body of the Ministry for Environment, Forest and Climate Change, Government of India under the signed agreement with certificate number IN—UK515716425337347W. This agreement for access and benefit sharing is Nagoya Protocol compliant, and we thank Dr. Hilke Marie Püschner for her assistance in procuring the certificate. *Anabaena* strain SSM-00 consortia was graciously provided by Dr. John Waterbury. Special thanks to Dr. Lars Möller for creating simplified images of host organisms based on electron micrographs and field photographs.

## Declarations

The authors declare that they have no competing interests.

## Supplemental Figures

**Fig. S1. Whole-genome visualization of newly circularized Peteryoungia and Hoeflea genomes.** Constructed using Circos (v0.69.8) within BV-BRC (v3.46.3), outside to inside tracks include contiguous sequence, navy blue; positive strand ORFs, green; negative strand ORFs, maroon; GC% content, black trace on pink zone; photosynthesis genes, blue. Chromosomes start with *dnaA* gene, whereas plasmids start with *repC*, marking locations in Mb or Kb. The concentric GC% content track illustrates absolute base composition across a 2,000-nt sliding window mapped on a fixed 0 to 1 scale, where the central dark grey dotted line denotes a baseline of 50% GC content and the surrounding thin grid lines indicate 25% and 75% thresholds inside to outside. Region with photosynthetic gene cluster (PGC) are enhanced as inserts to reveal limited GC% content shifts in each region, suggesting minimal genetic insertions/shifts. Note: Circular replicons not drawn to scale.

**Fig. S2. Photosynthetic operon structure and production of specific carotenoids. (A)** High gene synteny exists for all genes in photosynthetic gene cluster (PGC) operons of *Rhizobiaceae*, with a minor transversion in *H. anabaenae*, which matches deeper branching *Bradyrhizobium*. Genomes include (1) *P. albertimagni*, (2) *P. glycineendophyticum*, (3) *P. ipomoeae*, (4) *P. rhizophilum*, (5) ‘*Agrobacterium* sp. RAO6’, (6) *P. desertarenae*, (7) *P. rosettioformans*, (8) *P. aggregata*, (9) *H. olei*, (10) *H. phototrophica*, (11) *H. alexandrii*, (12) *H. anabaenae*, (13) *Bradyrhizobium.* sp. ORS 278, (14) *Bradyrhizobium* sp. BTAi1. *BChl* (green), *crt* (orange), *puf* (red), *puh* (pink), *hemA* (light blue), *lhaA/pucC* (purple), *acsF* (yellow), *regulator* (brown), *unbound cytochrome* (dark blue), *unknown gene* (white) depicted. LHII genes in *H. olei*, *H. phototrophica* (purple) match spectra (Fig. S5). *Bradyrhizobium* spp. further have RubisCo tandem enzymes to the puf operon (magenta), suggesting them to act as purple non-sulfur bacteria, rather than obligate anaerobic anoxgeninc phototrophs as for all *Hoeflea* and *Peteryoungia*. **(B)** As shown by gene content of PGCs, *Peteryoungia* spp. (P) and *Hoeflea anabaenae* (Ha) have the necessary genes for spirilloxanthin production, while all other *Hoeflea* (H) encode for spheroidene, further supported via the identification of such pigments via HPLC-MS/ HPLC-UV-VIS (Fig. S5).

**Fig. S3. Individual spectra of cultivated *Hoeflea* and *Peteryoungia* spp.** Whole cells (thick line) compared to pigment extract (thin line) reveals presence or absence of BChl *a* and carotenoids, which are bound in reaction centers and light harvesting complexes of specific strains. *Hoeflea* spp. generate more pigments under all conditions tested over *Peteryongia* spp, as seen by higher maxima of photosynthetic complexes in whole cells.

**Fig. S4. Influence of trace vitamins, elements, organics, and salinity on pigment production. (A)** PY medium supplemented with additional trace elements or vitamins reveals trace elements induce pigmentation in multiple strains. **(B)** PY medium supplemented with a gradient ASW (conditions 1 to 10 are 0%, 0.5%, 1%, 2%, 3%, 4%, 5%, 8%, 10%, 15% (w/v)) induces greatest pigmentation at 1-2% for both *Peteryoungia* and *Hoeflea* spp. **(C)** A gradient of complex organics (conditions 1 to 10 are 0%, 10%, 50%, 100%, 200%, 300%, 600%, 900%, 2000%, 3000% (w/v) std. PY medium) induces pigmentation at low concentration for both *Peteryoungia* and *Hoeflea* spp..

**Fig. S5. HPLC chromatograms of extracted pigments from photosynthetic *Rhizobiaceae*.** Carotenoid and bacteriochlorophyll compositions were determined by drying filtered extracts (0.2 µm pore size) under nitrogen gas flow prior to resuspension in 500 µL methanol:acetonitrile (85:15) mixed 10:1 with ammonium acetate (1 M), being run on a Kinetex C8 column and analyzed by LC-UV-VIS as compared to known spectra [**Nupur *et al.,* 2021**]. Predominant carotenoids and chlorophylls listed for each strain.

**Fig. S6. Core and shared genes of *Rhizobiaceae* members as compared to outgroups.** An upset plot shows the gene content shared among taxa, as compared to core genome tree previously generated. Core genes of class, family and genera listed in Table S5A, where unique genes of key species listed in Table S5B. *core genes shared between all members may be missing from genomes of draft status.

**Fig. S7. Photosynthesis gene cluster structure, evolution, and loss. (A)** Phylogenetic evolution comparison of 16S rRNA gene homology vs. concatinated photosynthesis genes *bchXZY-pufABC* of *Rhizobiales* among diverse Prokaryotic anoxygenic phototrophs. Congruent branching patterns suggests all larger groups such as the clade with *Hoeflea* and *Peteryoungia* (black arrow) to have contained photosynthesis for a long evolutionary period. Genera such as *Brevundimonas* or *Ectothiorhodospira* are not well resolved in the 16S rRNA gene tree, which inhibits the suggestion of vertical inheritance, as wel as suggestions of recent HGT of photosynthesis genes. **(B)** High gene synteny exists for all *Hoeflea* spp. as mapped with MAUVE alignment, showing identical locations of PGC (green box) in genome as compared to other genetic elements. **(C)** Comparing related phototrophic species *H. phototrophica, H. siderophila and H. olei* genomes reveals the lack of PGC, but retention of flanking genes (red boxes).

**Fig. S8. MLSA Tree of *Peteryoungia* spp. with additional high-quality MAGs available in databases.** The MLSA tree of 18 genomes compared via the AA sequence of 1000 concatenated genes has cultured-strains bolded, with sample (Symbiovar) source indicated as algal, light green; aquatic root, dark green; terrestrial roots, light orange; unknown, light grey. Photosynthesis genes present in bacteria, pink box; no photosynthesis genes, dark grey box; loss of photosynthesis at specific phylogenomic branches, red dots. Genomes utilized and core genes listed in Tables S7A & S7B.

**Fig. S9. MLSA Tree of *Hoeflea* spp. with additional high-quality MAGs available in databases.** The MLSA tree of 33 genomes compared via the AA sequence of 252 concatenated genes has cultured-strains bolded, with sample (Symbiovar) source indicated as algal, light green; terrestrial roots, light orange; unknown, light grey. Photosynthesis genes present in bacteria, pink box; no photosynthesis genes, dark grey box; loss of photosynthesis at specific phylogenomic branches, red dots. Genomes utilized and core genes listed in Tables S8A & S8B.

**Fig. S10. Ecological prevalence and potential niche of *Rhizobiaceae*.** Microbe atlas generated worldmaps of representatives *H. phototrophica*, *H. anabaenae*, *P. rosettiformans*, compared to *R. leguminosarum*, with individual dots representing samples with sufficient metadata to separate soil, aquatic, plant or animal samples, with sizes related to abundance. Inset pie-chart reveals proportions of total defined sample-types which contain each bacterial type.

## Supplemental Tables

**Table S1. Strains considered for growth experiments.** All type species of *Hoeflea* and related taxa, including *Peteryoungia* listed with isolation information, strain and genome availability, and published physiology.

**Table S2. Medium used for growth assays.** PY compared to modified PY+ medium composition used all strains to induce photosynthetic pigment production, along with ½SO medium modified for Anabaena growth.

Table S3. New circular genome info of *Hoeflea* and *Peteryoungia* spp. Genome information included for chromosome and plasmids.

**Table S4. *Rhizobiales*-wide presence/absence of nodulation, photosynthesis, and nitrogen fixation genes.** 7527 high quality genomes including our newly sequenced species and all Rhizobiales of GTDB filtered by BacRep were screened for genes of interest by a HMMER3 search of protein families among each genome, depicting presence of each gene with a “1”.

**Table S5. *Rhizobiaceae* high-quality genomes utilized for gene content differentiation among newly sequenced *Hoeflea* and *Peteryoungia*. (A)** Accession numbers and genes associated to photosynthesis, nitrogen-fixation, nodulation, symbiosis/pathogenesis, vitamin synthesis, and iron acquisition of the 77 genomes used for the MLSA genome tree (Fig. 3), which utilized **(B)** 208 single copy core genes that occurred in all *Rhizobiales* members. **(C)** The core genes of chosen *Rhizobiales*, *Rhizobiaceae*, or genera as calculated and represented in the upset plot (Fig. S6), with **(D)** genes unique specific key species also reported.

**Table S6. *Rhizobiaceae* ancestral genome gain, loss and duplication events. (A)** Orthologous gene groups identified in Orthofinder utilized for GLADE evolutionary analysis to determine **(B)** core genes per ancestral node, **(C)** gene gains per node, **(D)** genes lost per node, and **(E)** genes lost per node.

**Table S7. *Peteryoungia* metagenomes utilized in comparative genomics. (A)** Accession and related information including source of high-quality MAGs for the 18 genomes used for the MLSA tree (Fig. S8), which utilized **(B)** 1000 single copy genes representing PGFams.

**Table S8. *Hoeflea* metagenomes utilized in comparative genomics. (A)** Accession and related information including source of high-quality MAGs for the 34 genomes used for the MLSA tree (Fig. S9), which utilized **(B)** 252 single copy genes representing PGFams.

**Table S9. *Prokaryotic* genomes utilized for the evolutionary time-tree analysis. (A)** The 20 hmm ribosomal gene references, accession numbers and related information for the 181 genomes for the MLSA tree utilized for the time-scale evolutionary history analysis (Fig. 5). **(B)** The 12-fossil record calibration timepoints/ events chosen to strap the phylogenetic tree to the ancestral record (Fig. 5).

**Table S10. Cultivation-independent sequence-based ecology as determined by Microbe-atlas 16S geography analysis. (A)** *Rhizobium leguminosarum* database search compared to *Hoeflea phototrophica*, *Hoeflea anabaenae*, *Peteryoungia aggregatum* and *Peteryoungia rosettiformans* reveals overall environmental habitats, whereas **(B)** top 3 sample hits of highest density are listed per species suggesting specific ecological niche.

## Notes

### Competing Interest Statement

The authors have declared no competing interest.

