## Supplemental Figures for "Ancestral phototrophic *Rhizobiaceae* evolved in association with algae, then plants"

**Fig. S10. Ecological prevalence and potential niche of *Rhizobiaceae*.** Microbe atlas generated worldmaps of representatives *H. phototrophica*, *H. anabaenae*, *P. rosettiformans*, compared to *R. leguminosarum*, with individual dots representing samples with sufficient metadata to separate soil, aquatic, plant or animal samples, with sizes related to abundance. Inset pie-chart reveals proportions of total defined sample-types which contain each bacterial type.

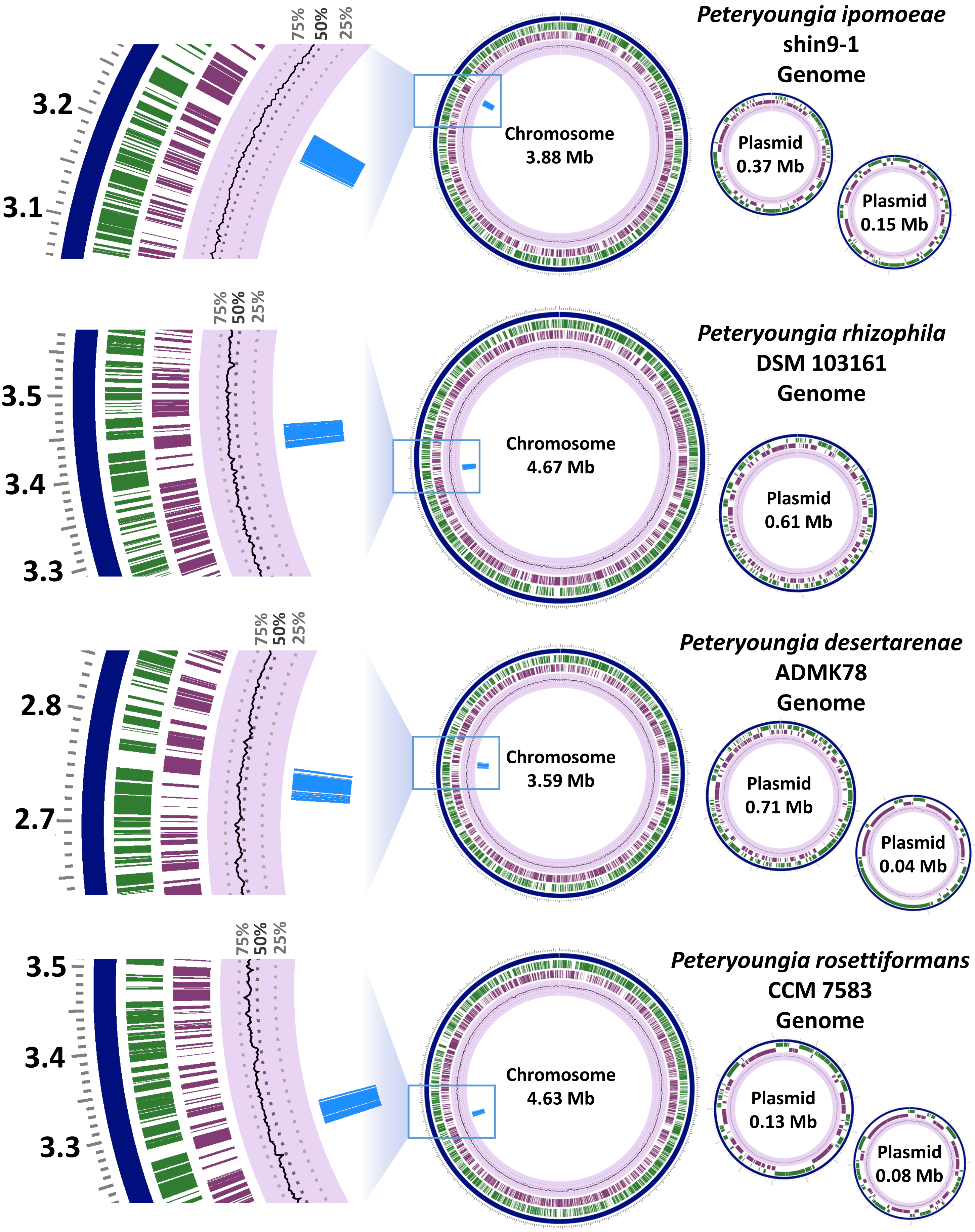


**Fig. S1. Whole-genome visualization of newly circularized *Peteryoungia* and *Hoeflea* genomes.** Page 1/3.

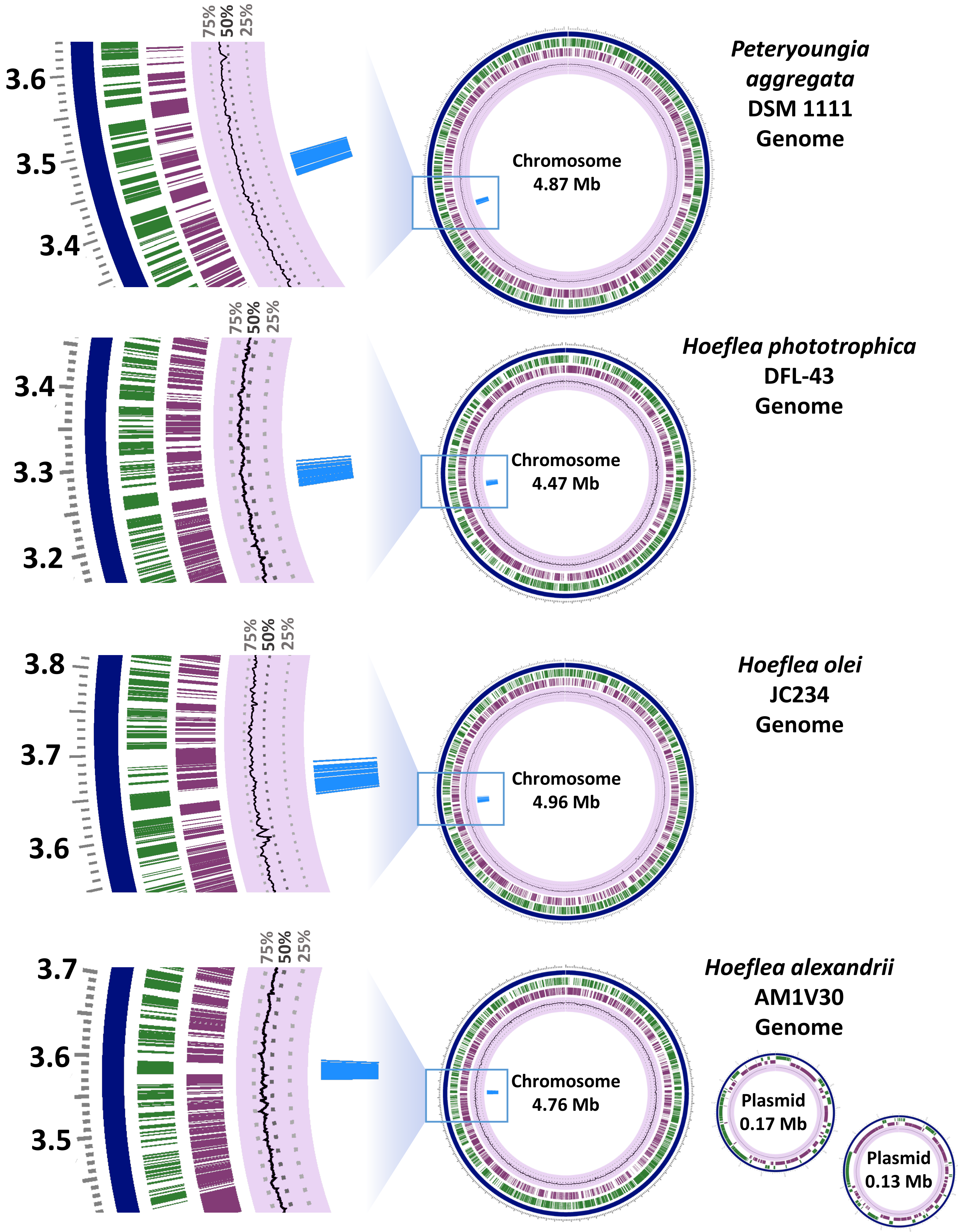


**Fig. S1. Whole-genome visualization of newly circularized *Peteryoungia* and *Hoeflea* genomes.** Page 2/3.

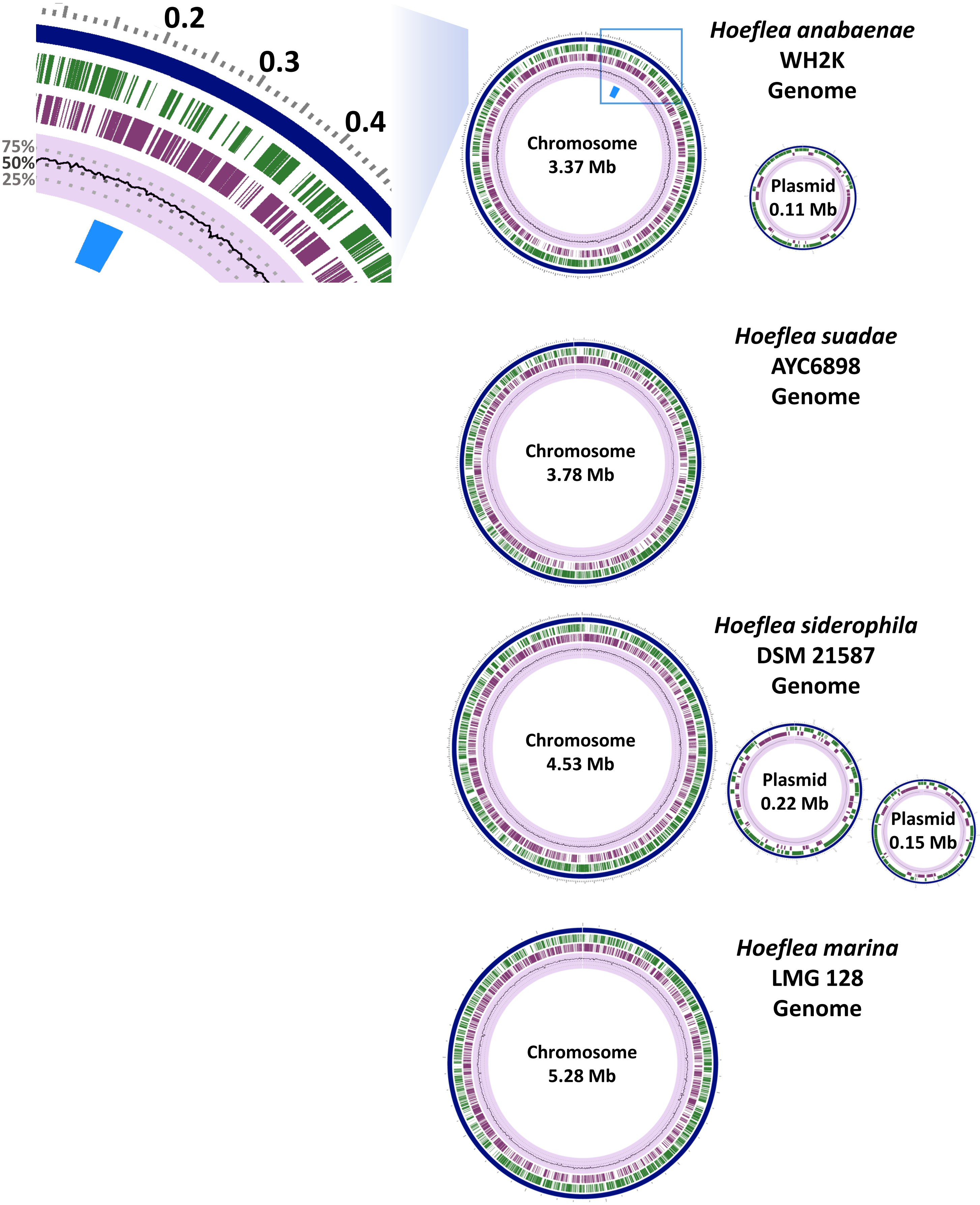


**Fig. S1. Whole-genome visualization of newly circularized *Peteryoungia* and *Hoeflea* genomes.** Page 3/3.

A

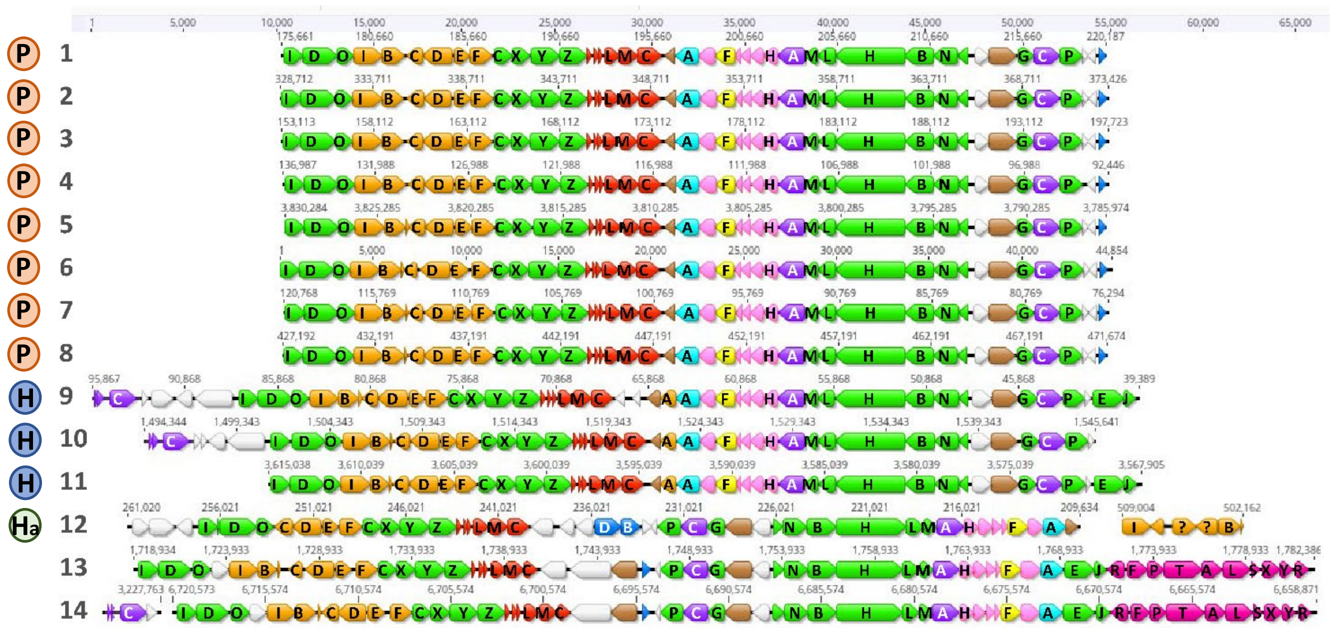

B

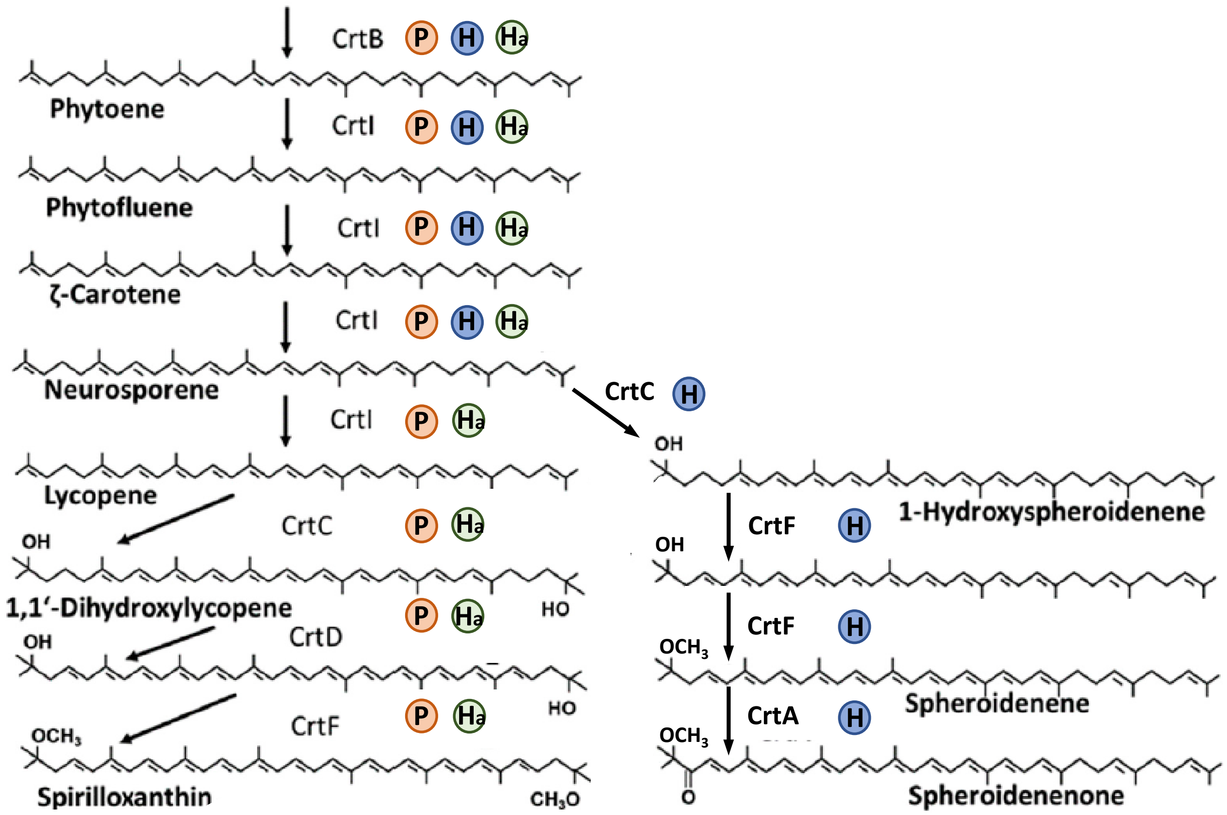


**Fig. S2. Photosynthetic operon structure and production of specific carotenoids.**

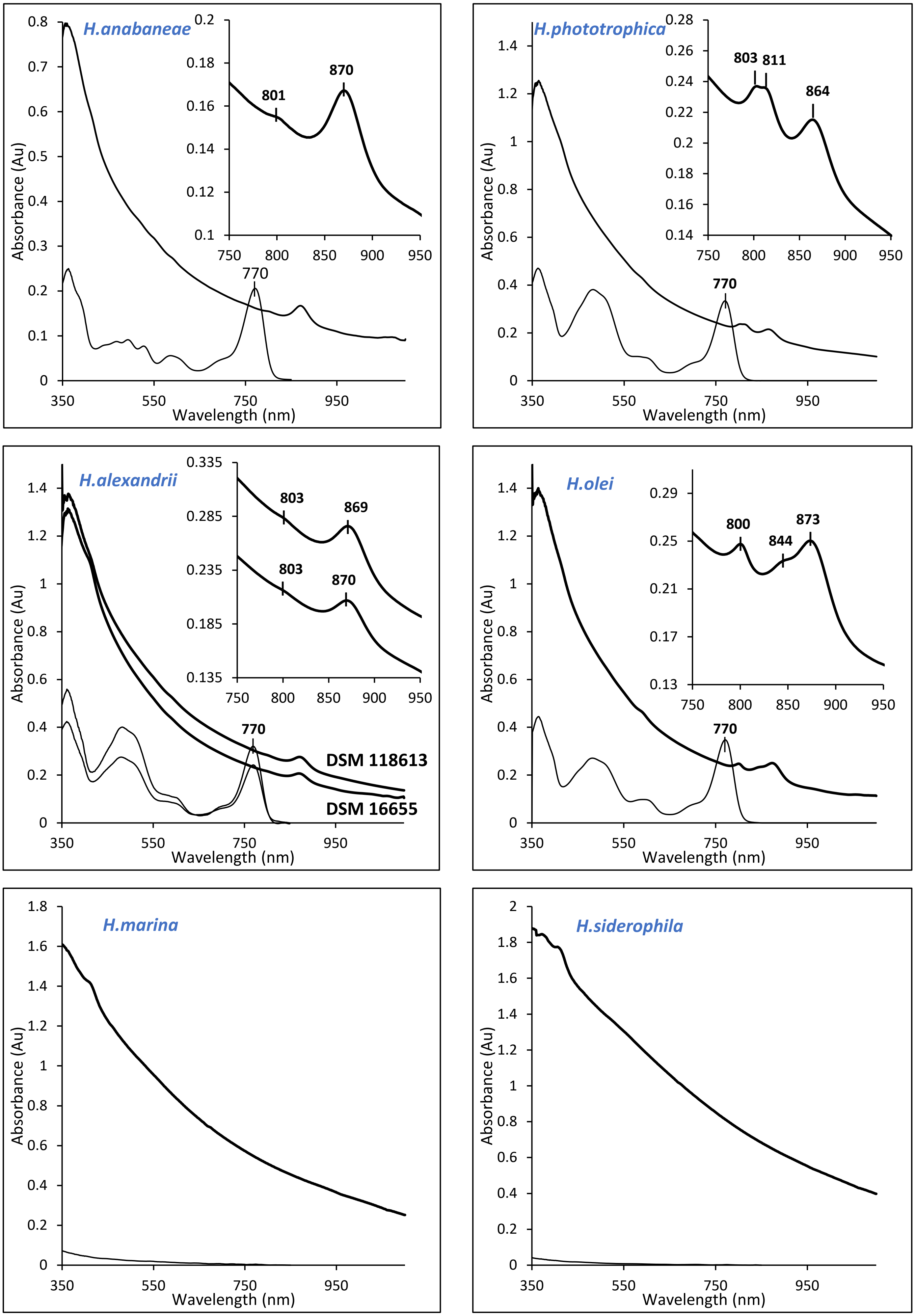


**Fig. S3. Individual spectra of cultivated *Hoeflea* and *Peteryoungia* spp.**

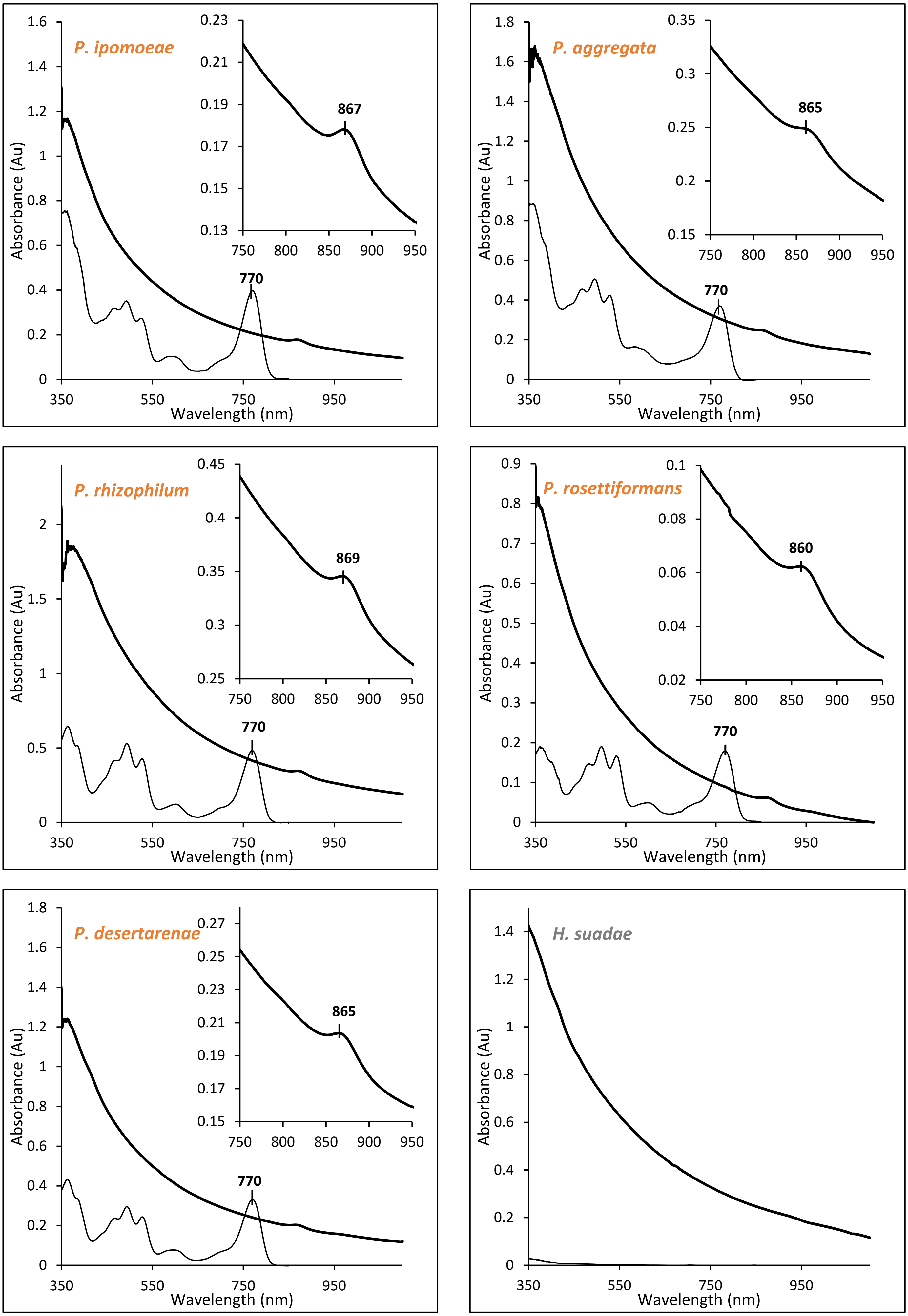


**Fig. S3. Cont. Individual spectra of cultivated *Hoeflea* and *Peteryoungia* spp.**

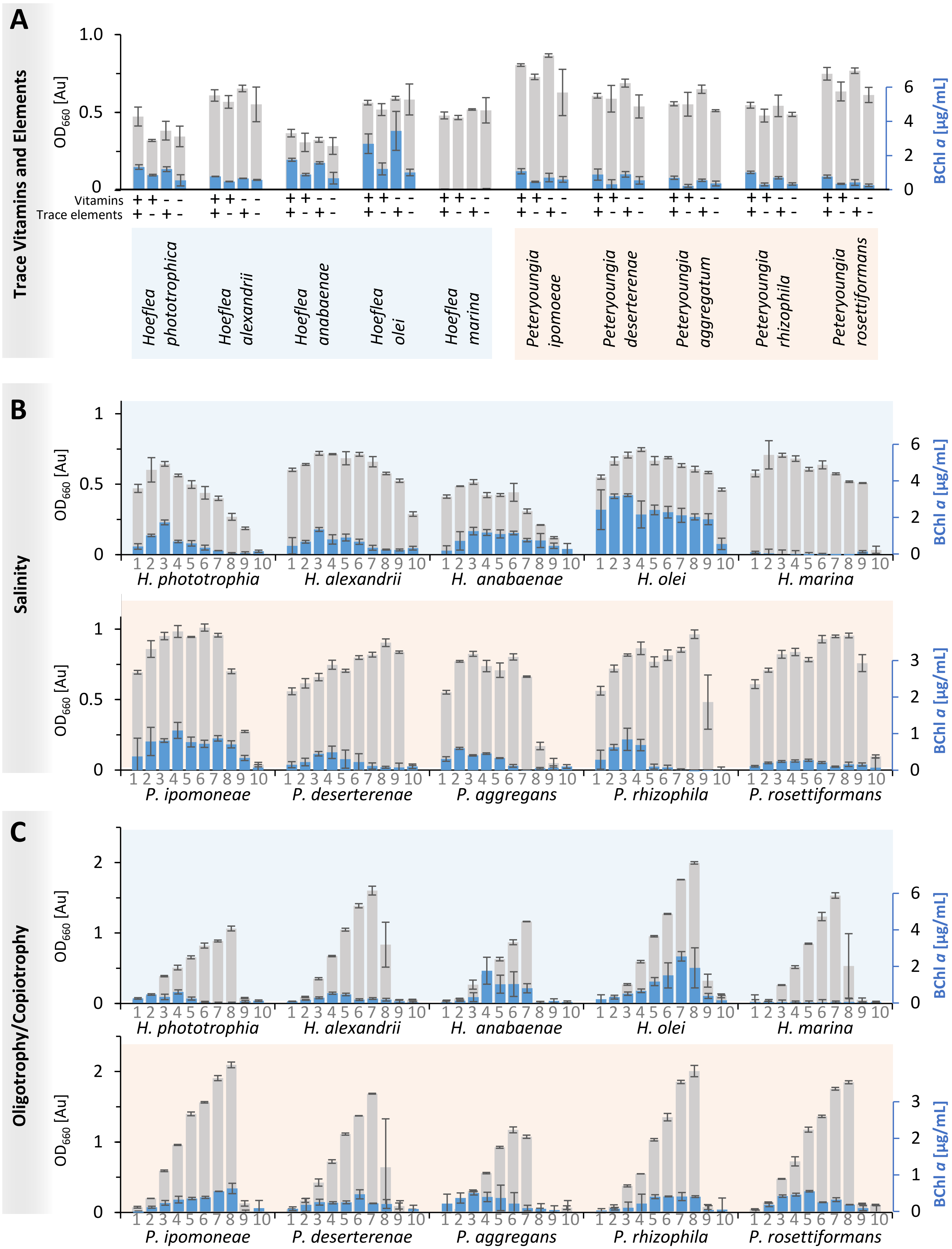


**Fig. S4. Influence of trace vitamins, elements, salinity, and organics on pigment production.**

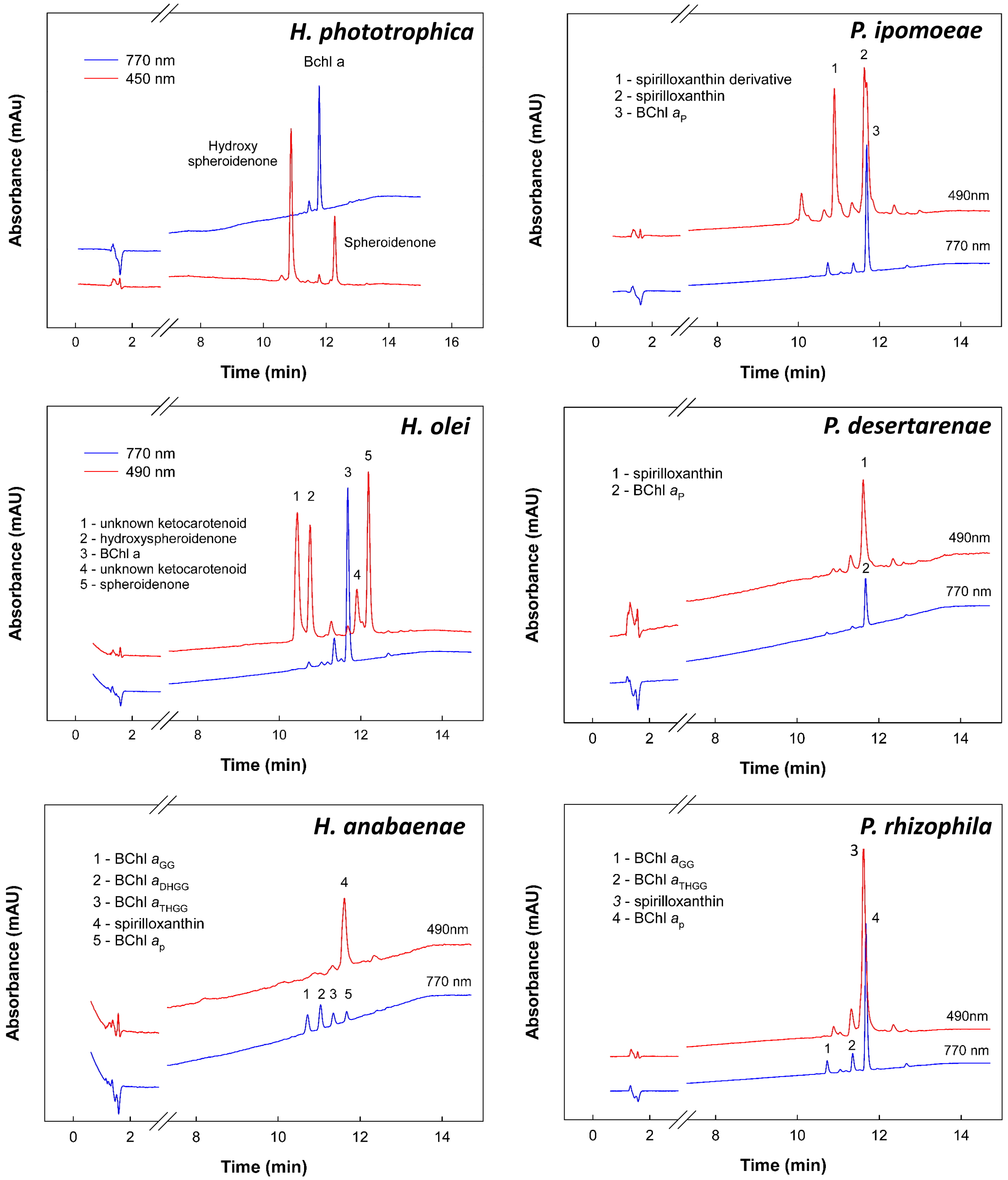


**Fig. S5. HPLC chromatograms of extracted pigments from photosynthetic *Rhizobiaceae*.**

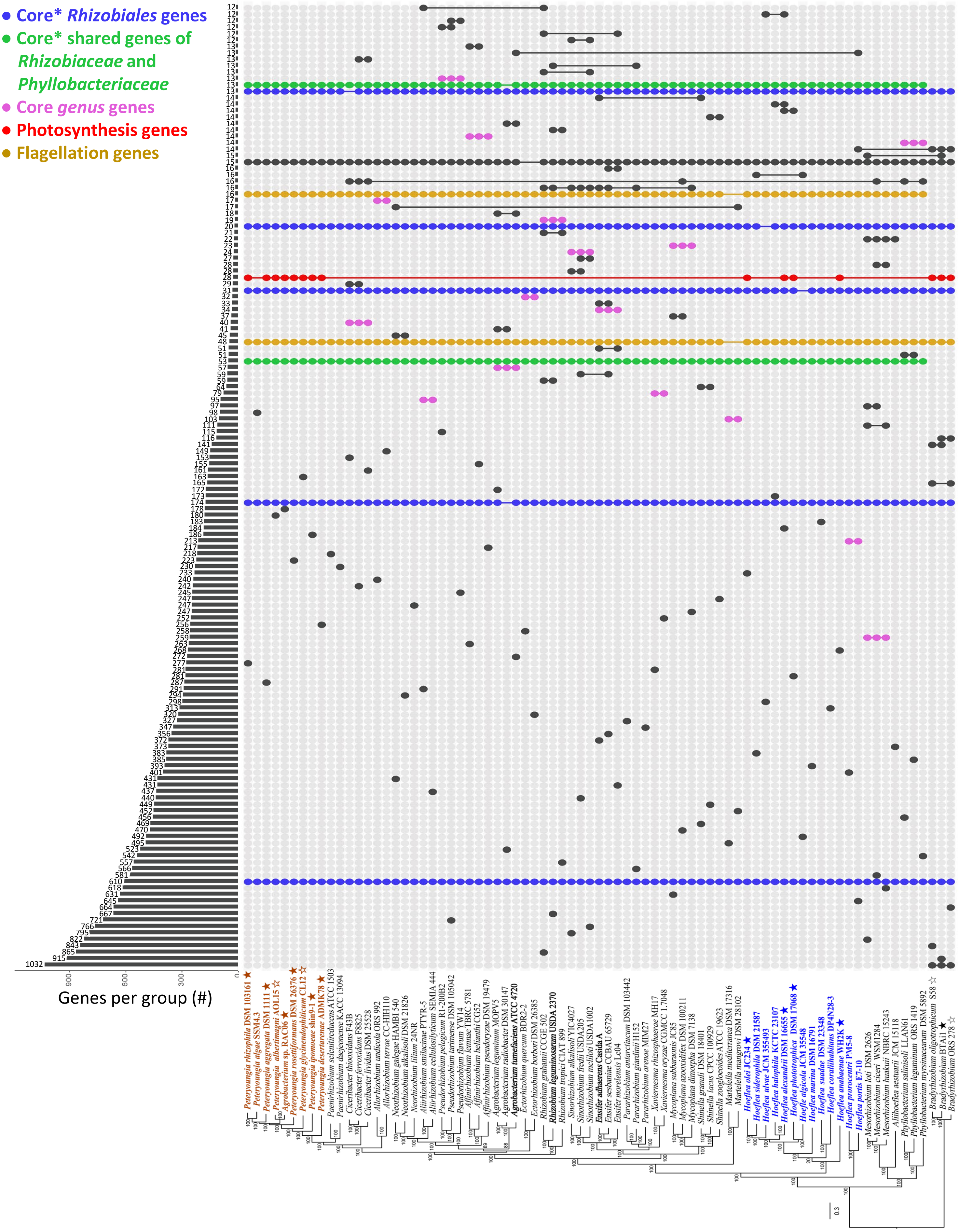


**Fig. S6. Core and shared genes of *Rhizobiaceae* members.**

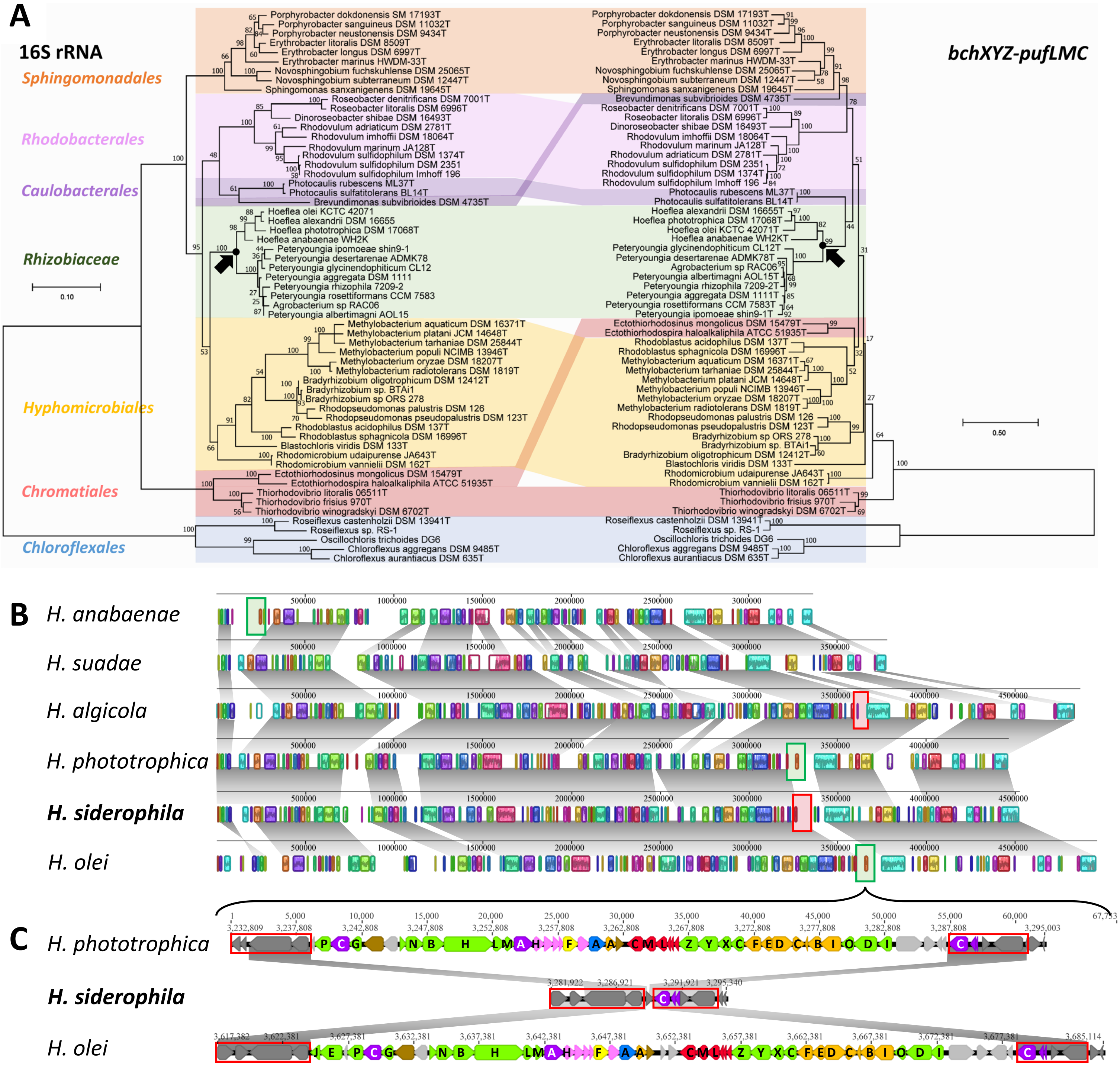


**Fig. S7. Comparative genomics of *Rhizobiaceae* members reveals vertical inheritance of photosynthesis, and successive and gene loss.**

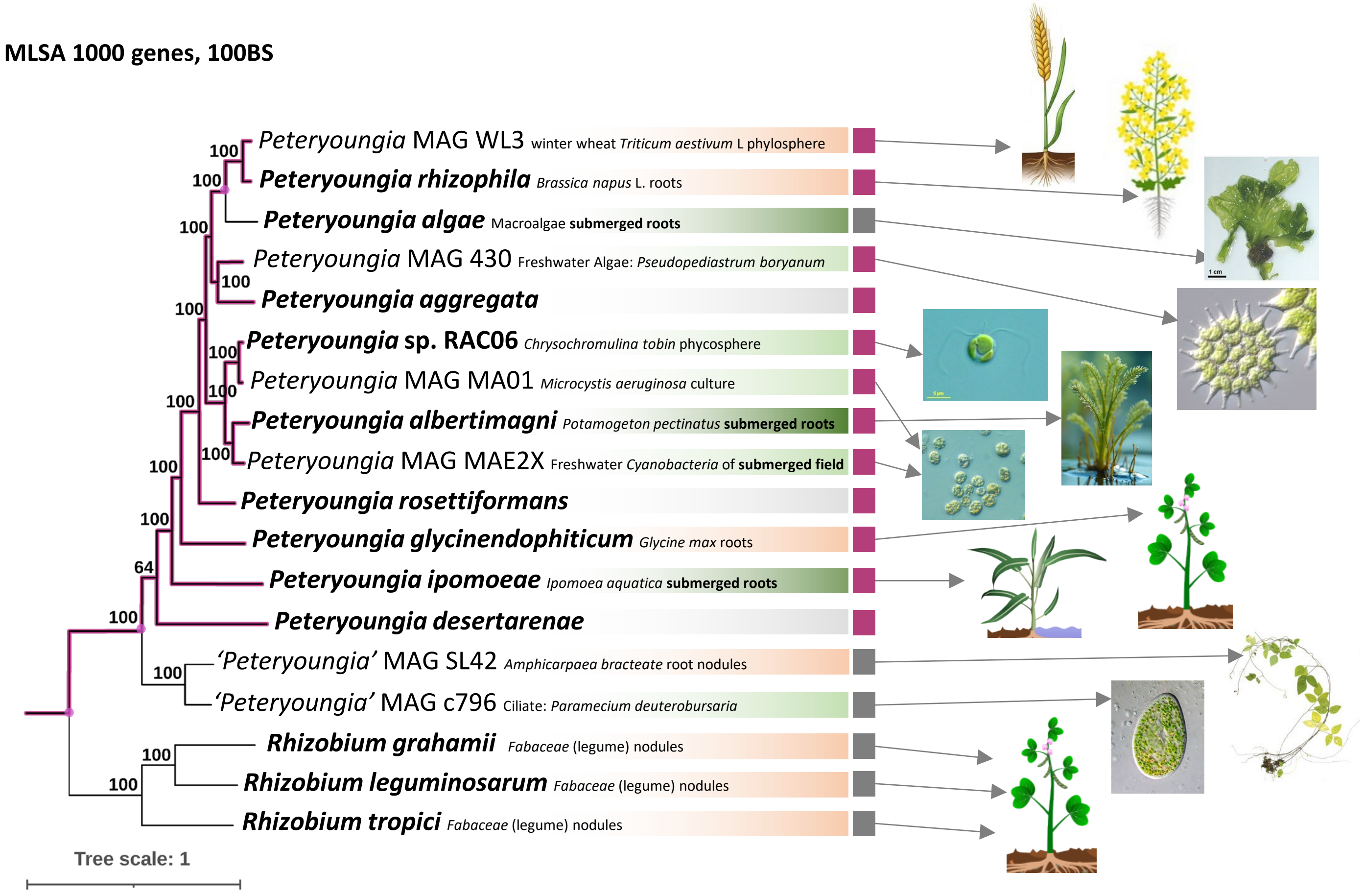


**Fig. S8. MLSA Tree of *Peteryoungia* spp. with additional high-quality MAGs available in databases.**

MLSA 252 genes, 100BS

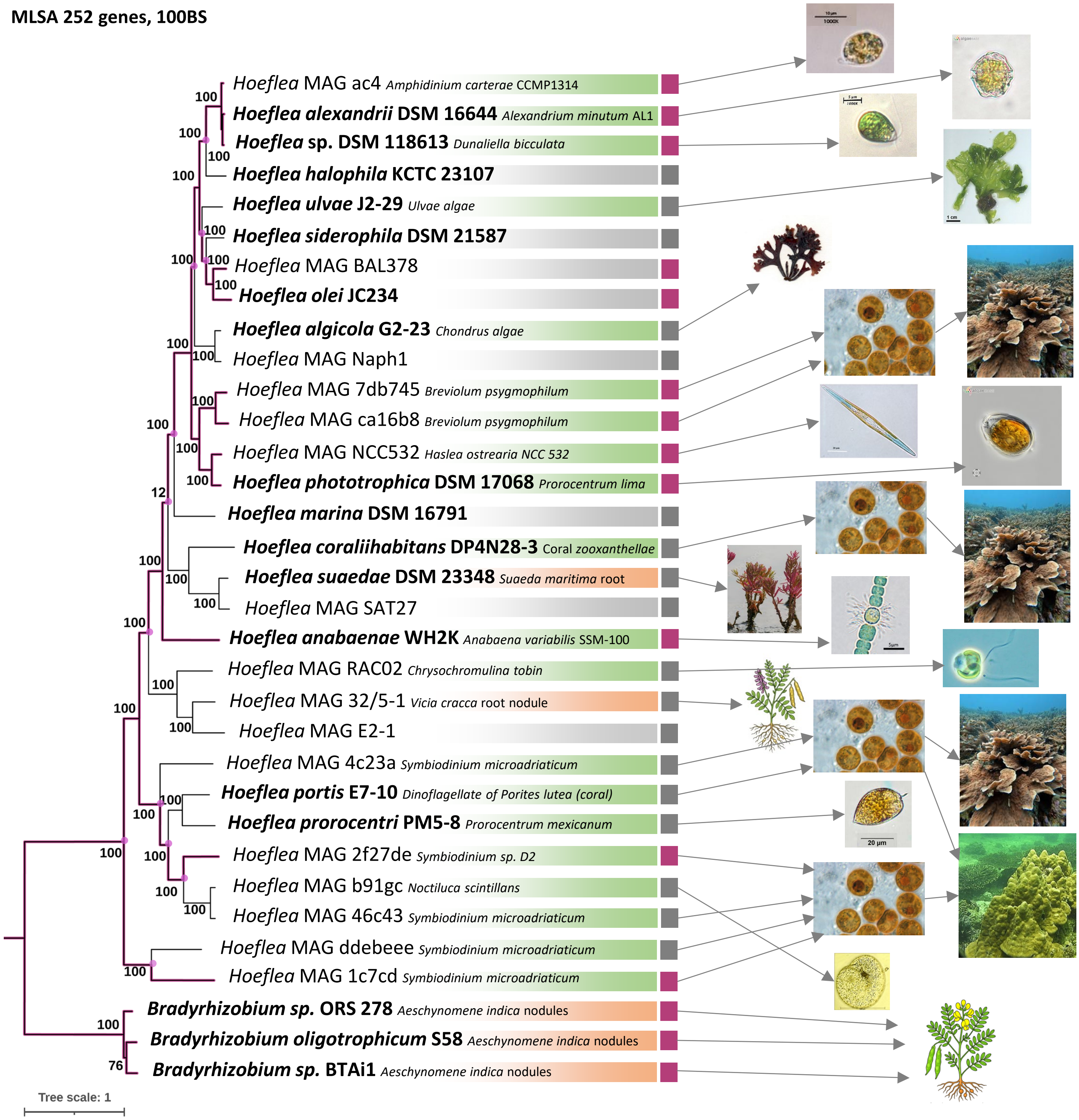


**Fig. S9. MLSA Tree of '*Hoeflea*' spp. with additional high-quality MAGs available in databases.**

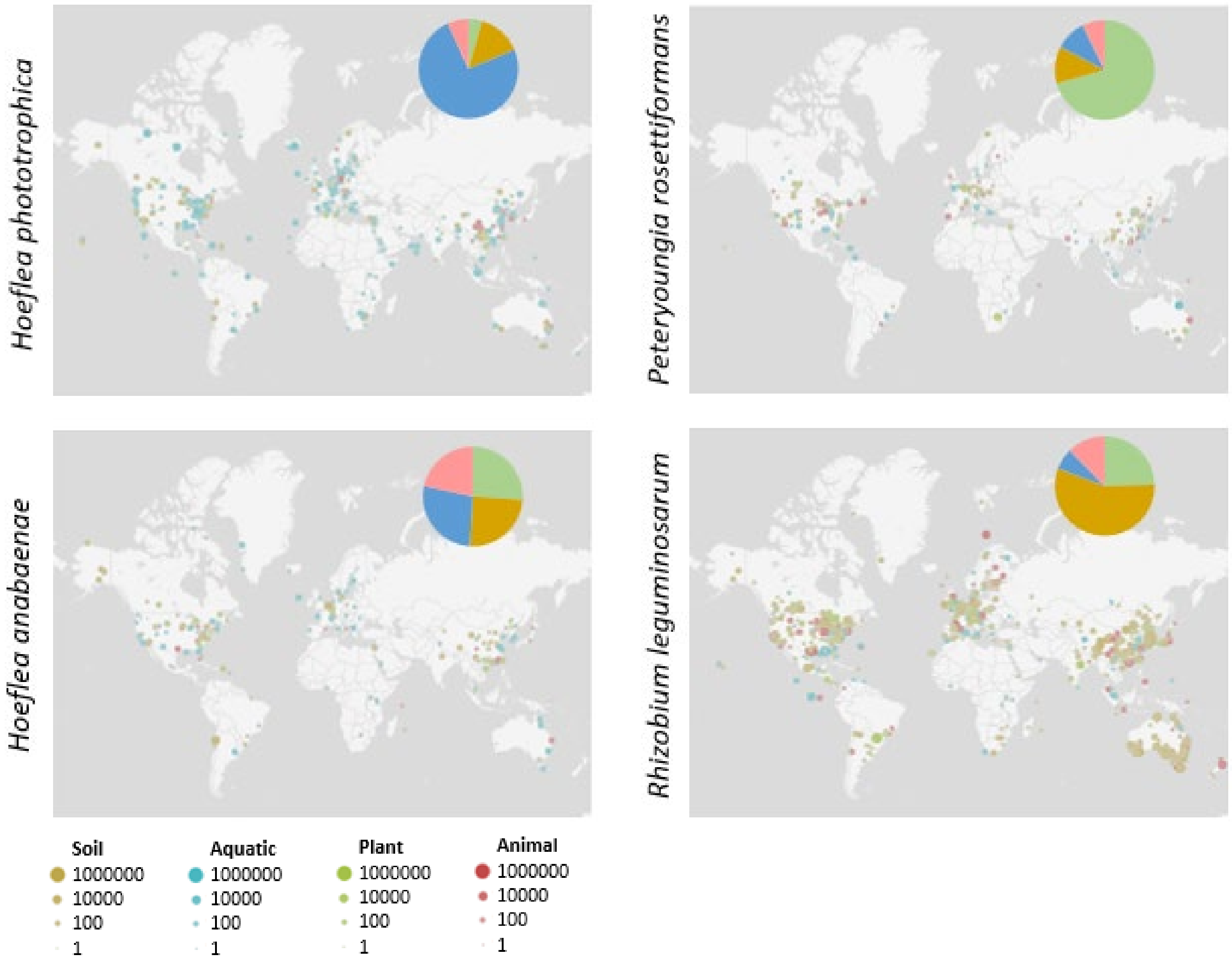


**Fig. S10. Ecological prevalence and potential niche of *Rhizobiaceae*.**
